# Extracellular vesicles and co-isolated endogenous retroviruses differently affect dendritic cells

**DOI:** 10.1101/2023.01.27.525863

**Authors:** Federico Cocozza, Lorena Martin-Jaular, Lien Lippens, Aurelie Di Cicco, Yago A Arribas, Florent Dingli, Michael Richard, Louise Merle, Patrick Poullet, Damarys Loew, Daniel Lévy, An Hendrix, George Kassiotis, Alain Joliot, Mercedes Tkach, Clotilde Théry

**Affiliations:** INSERM U932, Institut Curie Centre de Recherche, PSL Research University, F-75005, Paris, France; Université de Paris, F-75006, Paris, France; Institut Curie centre de recherche, CurieCoreTech Extracellular Vesicles, F-75005, Paris, France; Laboratory of Experimental Cancer Research, Ghent University, Ghent, Belgium; Institut Curie, PSL Research University, Sorbonne Université, CNRS UMR168, Laboratoire Physico-chimie Curie, F-75005, Paris, France; Institut Curie, PSL Research University, CNRS UMR144, Cell and Tissue Imaging Facility (PICT-IBiSA), F-75005, Paris, France; Institut Curie, PSL Research University, Centre de Recherche, CurieCoreTech Spectrométrie de Masse Protéomique, F-75005, Paris, France; Institut Curie, Bioinformatics core facility (CUBIC), INSERM U900, PSL Research University, Mines Paris Tech, F-75005, Paris, France; Retroviral Immunology, The Francis Crick Institute and Department of Medicine, Faculty of Medicine, Imperial College, London UK

**Keywords:** antigen presenting cells, exosomes, extracellular vesicles, tumors, retrovirus

## Abstract

Cells secrete membrane-enclosed extracellular vesicles (EVs) and non-vesicular nanoparticles (ENPs) that may play a role in intercellular communication. Tumor-derived EVs have been proposed either to induce immune priming of antigen presenting cells, or, to be immuno-suppressive agents promoting tumor immune escape. We suspect that such disparate functions are due to variable composition in EV subtypes and ENPs of the analyzed EV preparations. We aimed to exhaustively characterize the array of secreted EVs and ENPs of murine tumor cell lines. Unexpectedly, we identified virus-like particles (VLPs) from endogenous murine leukemia virus in preparations of EVs produced by tumor cells. We established a robust protocol to separate small (s)EVs from VLPs and ENPs. We compared their protein composition and analyzed their functional interaction with target dendritic cells (DCs). ENPs were poorly captured and did not affect DCs. sEVs specifically induced DC death. A mixed EV/VLP preparation was the most efficient to induce DC maturation and antigen presentation. Our results call for systematic re-evaluation of the respective proportions and functions of non-viral EVs and VLPs produced by tumors and their contribution to anti-tumor immune responses and to tumor progression.

## INTRODUCTION

Intercellular communication is crucial in all tissues, and especially in the tumor microenvironment, where cancer cells must instruct the surrounding stromal and immune cells to eventually form a tumor. Cells can interact with each other and with the extracellular medium through soluble secreted factors, but also through extracellular vesicles (EVs) and other non-vesicular extracellular nano-particles (ENPs). In the last years, the development of novel isolation and detection technologies of EVs have increased our knowledge on EVs/ENPs heterogeneity (Tkach et al., 2018; Willms et al., 2018), and highlighted the existence of novel subtypes of non-vesicular ENPs such as exomeres (Zhang et al., 2018), supermeres (Zhang et al., 2021b), and supramolecular attack particles (Balint et al., 2020). However, except for the latter which display killing activity, little is known still on the biological functions of these recently discovered particles (Zhang et al., 2019). In addition, most of the functional capacities attributed to EVs were observed with a mixture of EVs subtypes, ENPs and other components collectively considered as contaminants (soluble proteins, lipoproteins, etc.), raising the questions of the specific properties of each subtype of EV and ENP.

Viral infection brings an additional level of complexity to the vesicular secretome. Viruses depend on biological processes of the host cells to survive and to form viral particles. In particular, enveloped viruses are enclosed in a host cell-derived lipid bilayer which, for instance in the case of HIV, uses the EV formation machinery to be formed (Booth et al., 2006; Gould et al., 2003). Therefore, infection by viruses modifies the amount and composition of the EVs, which can then also include viral components (Martin-Jaular et al., 2021; Nolte-’t Hoen et al., 2016). Endogenous retroviruses (ERVs) represent a potential source of enveloped viral particles and of modified EVs. ERVs are inserted in the genome of higher organisms, but are not necessarily part of a pathologic process, since they are normally mutated or repressed and thus cannot form infectious viral particles (Kassiotis and Stoye, 2017). However, their infectivity can be restored in physiological or pathological conditions, for instance in immunodeficient models or in mouse tumors (Ottina et al., 2018; Young et al., 2012).

In the tumor microenvironment, tumor cells promote changes in surrounding cells of the stroma and the immune system (Gajewski et al., 2013; Whiteside, 2008), in part through the action of EVs and ENPs (Han et al., 2019). However, the actual effects of EVs/ENPs released by tumor cells towards the immune system are controversial: there is evidence showing both immune response activation resulting from tumor antigen transfer, and conversely, immunosuppressive functions (Greening et al., 2015; Robbins and Morelli, 2014; Thery et al., 2009). This discrepancy could be due to the different isolation methods and conditions used in these studies, leading to different combinations of heterogenous EVs, ENPs and associated contaminants, each bearing their specific and sometimes antagonist functions.

Tumor-derived EVs (endogenous or by bio-engineering) can carry antigens from the producing cells, either native or as peptides presented in MHC-I complexes on the EV membrane, and transfer it to dendritic cells (DCs), leading to a T cell-mediated antitumor response and tumor rejection (Chulpanova et al., 2018; Wolfers et al., 2001; Zeelenberg et al., 2008). Immature DCs surveil the peripheral tissues and have a high uptake capacity. Once they have captured antigens and sensed the microenvironment, they can eventually mature and migrate to lymph nodes, especially if they have detected damage, pathogen or inflammatory molecules. There, they can activate or tolerize T cells, depending on the sensed signals (Banchereau and Steinman, 1998). In addition to containing antigens, it has been proposed that EVs possess an adjuvant-intrinsic effect due to the presence of damage-associate molecular patterns (DAMPs), which may explain why EVs are more efficient than soluble proteins at transferring antigens to antigen-presenting cells (APCs) (Morelli et al., 2004; Robbins and Morelli, 2014). However, whether the antigen transfer capacities or the DAMPs are carried by all EV subtypes or ENPs, or only some specific types is unknown.

Here, we have tackled this question by first characterizing the heterogeneity of EVs and ENPs isolated from the murine mammary tumor cell line EO771. Surprisingly, we found a remarkable amount of enveloped virus-like particles in our preparations, that we identified as infectious retroviruses produced from endogenous murine leukemia virus (MLV). This was also observed in EVs from two other murine breast (4T1) or dendritic (MutuDC) tumor cell lines, but not in primary DCs nor immortalized fibroblastic cells. We established a protocol to reliably separate the VLPs from other small EVs and ENPs. We characterized the tumor particle subtypes by quantitative proteomic analysis and analyzed their respective capacity to transfer antigen and activate dendritic cells in vitro. Unexpectedly, our results unravel a limited potential of VLP-devoid small EVs for the induction of antigen-specific immune responses. By contrast, the large/small/VLP mixed EVs subtypes displayed the highest immune activation potential.

## RESULTS

### EO771 cells secrete heterogenous particles including EVs, ENPs and VLPs into the extracellular medium

We chose the EO771 mouse cell line as a model of breast tumor cells that can be used for downstream analysis of antigen presentation in the syngeneic mouse C57BL/6 genetic background. To explore the heterogeneity of the EVs and particles secreted by EO771, we first performed a rough separation of large/medium EVs, small EVs and ENPs, based on classical differential centrifugation protocols (Jeppesen et al., 2019; Thery et al., 2006). Briefly, the serum-free conditioned medium depleted of dead cells and largest particles by a 2,000xg centrifugation was concentrated using 10 kDa filters, followed by sequential ultracentrifugation at 10,000x*g* (16 min), 200,000x*g* (50 min) and 200,000x*g* overnight (ON) to obtain pellets called 10k, 200k and ENPs respectively (Fig.1A). Nanoparticle tracking analysis (Fig.1B) showed that, as expected, the 10k pellets contained more EVs larger than 200 nm in diameter than the 200k and ENPs. Of note, however, by NTA analysis, all pellets contained a majority of small EVs (< 200 nm).

**Figure 1:**
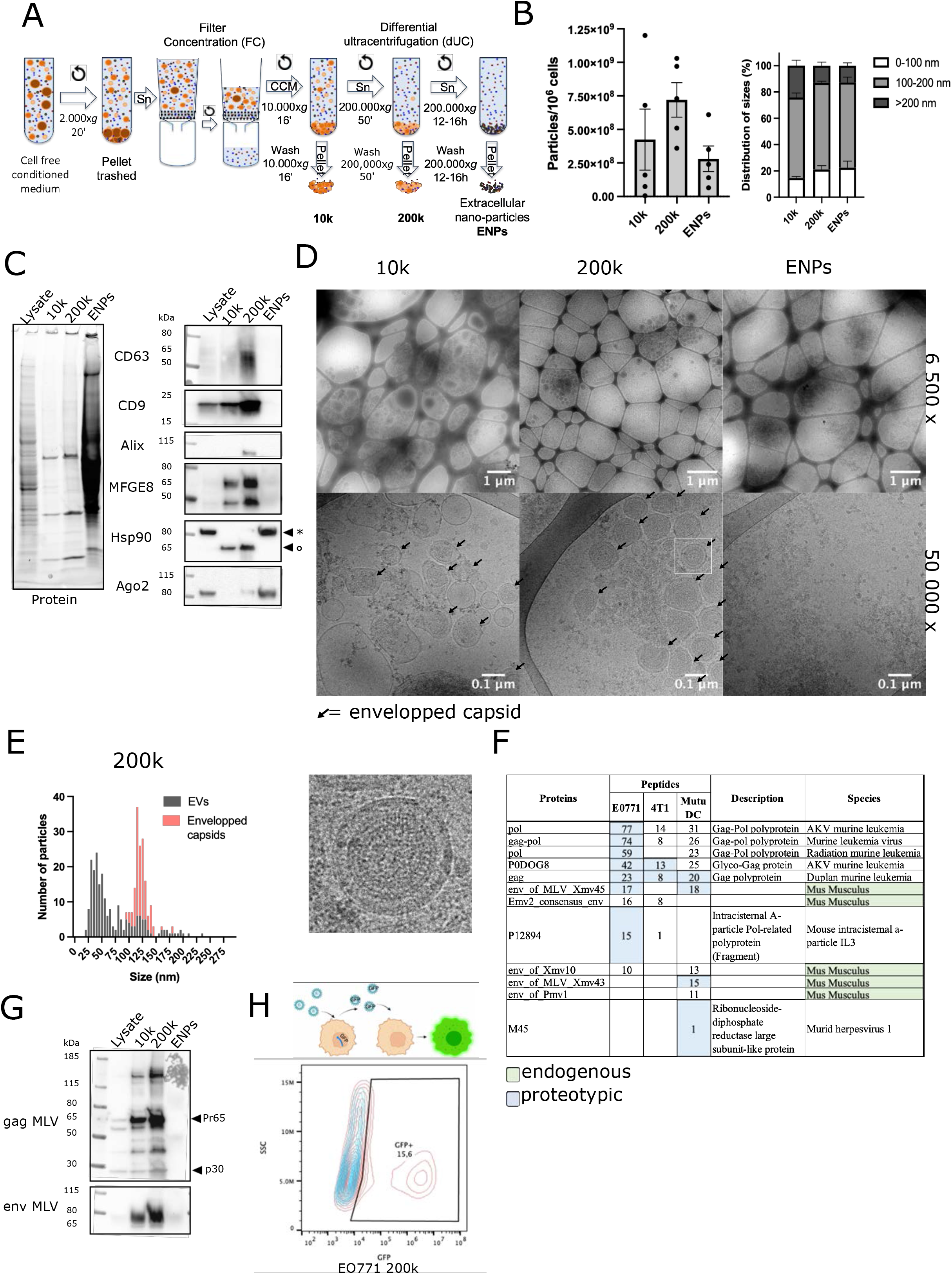
Characterization of the EVs and ENPs from EO771 cell line: presence of virus-like particles (VLPs). (A) Scheme of the protocol of separation of 10k, 200k and ENPs. (B) NTA analysis of 10k, 200k and ENPs: particle quantification (left) and size distribution (right) in n=4. (C) Western blot showing total proteins (left) and various protein markers as indicated (right). Lysate from 2×10^5^ EO771 cells, 10k, 200k and ENPs secreted by 20×10^6^ EO771 cells were loaded on the gel. * indicates specific Hsp90 band of the expected size. ° indicates a shorter size band resulting from endogenous cleavage. (D) Cryo-EM of EO771 10k, 200k and ENPs. Arrows indicate typical capsid-like structures inside EVs. One such structure highlighted by a white square is shown in a zoom (below, right). (E) Quantification of size and presence (“enveloped capsids”) or absence (EVs) of a VLP structure in all particles of one representative sample of 200k analyzed by cryo-EM (from images at 50,000X magnification). (F) Proteomic identification of viral proteins in 200k pellets of EO771, 4T1 and MutuDC. Blue: proteins identified by specific proteotypic peptides. Green: proteins from our curated endogenous retrovirus database. (G) Western blot shown in C) incubated with antibodies anti-MLV proteins gag and env. (H) Infectivity capacity of the 200k from EO771 cells, shown by production of pseudotyped XG7 GFP-encoding virus from *Mus Dunni-*XG7 cells, 14 days after exposure to the EO771 200k. Pseudotyped virus in the supernatant of *Mus Dunni*-XG7 was evidenced by detection of GFP expression in parental *Mus Dunni* exposed to this supernatant (red), as compared to cells not exposed to the supernatant (blue).

The 10k, 200k and ENP pellets obtained from the same number of producing cells (20×10^6^) were run on a western blot together with the lysate of the producing cells (2×10^5^) and revealed with antibodies against transmembrane (CD9, CD63), external membrane-bound (Milk-fat globule-EGF-factor VIII, MFGE8), and cytosolic proteins (Alix, Ago2, Hsp90) classically used to characterize EVs and/or ENPs (Fig.1C). The total protein stain showed that ENPs contained the highest amount of proteins, followed by the 200k and 10k. However, looking at specific markers, the 200k was enriched in CD63, CD9 and MFGE8, also found in the 10k but not in the ENPs. By contrast, Alix was only detected in the 200k. Conversely, Ago2 was mainly detected in the ENPs, and to a lesser extent in the 200k. Finally, the full-length Hsp90 (*, Fig.1C) was found only in the cell lysate and the ENPs but not in the 10k or 200k, although a shorter band of around 70kDa (°, Fig.1C) was detected in the latter, probably corresponding to a previously described stress-induced cleaved form (Beck et al., 2009).

Cryo-electron microscopy (cryo-EM) (Fig.1D) revealed a great heterogeneity of particles in the 10k and 200k, including vesicles of various sizes and aspects, as well as smaller particles without any apparent lipid bilayer and aggregates of electrodense material. ENPs contained mainly the latter two structures. Surprisingly, some of the vesicles in the 10k and 200k contained circular structures with regular concentric strips (arrows and close-up in Fig.1D), a shape typical of viral capsids (Qu et al., 2018), suggesting the presence of virus-like particles (VLPs). We quantified the size and the VLP aspect of the vesicles in the 200k (Fig.1E). VLP-shaped vesicles were concentrated in the range of 100-150 nm diameter (peak at 115-130 nm), consistent with the observed size of in vitro reconstituted murine leukemia virus (MLV) (Qu et al., 2018), but never observed in the smaller vesicles (25-60 up to 80 nm diameter).

To determine whether these particles were actually of viral origin, we performed a qualitative mass spectrometry-based proteomic analysis of the 200k. Mass spectrometry files were interrogated, to identify peptides from these proteins, against a data base containing the mouse proteome concatenated to protein sequences from all known mouse viruses (523 sequences manually extracted from Swissprot) and from 53 sequences of endogenous MLV envelope glycoproteins translated from the nucleotide sequences of proviruses annotated as previously described (Attig et al., 2017) (Fig.1F, Suppl. Table 1). We evidenced the presence of several proteins of the family of murine leukemia virus (MLV), both exogenous (AKV, Duplan) and endogenous (Xmv, Intracisternal A-particle). This was not specific of the EO771 EVs, since we also identified mouse retroviral proteins (Fig.1F,Suppl. Fig.1B, Suppl. Table 1) and VLP structures (Suppl. Fig.1A) in 200k pellets of another breast carcinoma cell line, 4T1 (Balb/c genetic background), and in a tumor dendritic cell line (MutuDC, C57BL/6 background). By contrast, we did not see VLP structures by cryo-EM in 200k samples from primary bone marrow-derived DCs (BMDCs) (Suppl. Fig.1A).

The western blot shown in Fig.1C was further probed with antibodies against gag and envelope (env) proteins from MLV (R187 and 83A25 antibodies) (Fig.1G), which revealed the presence of both proteins in the 200k, and to a lesser extent in the 10k, but not in the ENPs. Gag and env were also detected in the 200k from 4T1 and MutuDC (Suppl. Fig.1B), and only env (but not gag) was detected in EVs from a spontaneously immortalized fibroblastic cell line, Pfa1 (Suppl. Fig.1B). Finally, we evaluated the infectivity capacity of these viral particles by exposing permissive *Mus Dunni*-XG7 cells (where a GFP-encoding reporter provirus is mobilized upon retroviral infection) to the 200k of EO771 for 24h. The cell-conditioned medium (CCM) was then collected once per week during 2 weeks, and the last CCM was added to unmodified *Mus Dunni* (Young et al., 2012): detection of GFP-expressing cells in the CCM-exposed *Mus Dunni* cells confirmed the presence of infectious retroviral particles in the 200k of EO771 (Fig.1H), and of MutuDCs (Suppl. Fig1C). Therefore, the VLP preparations also contain infectious retroviruses. However, not knowing the proportions of infectious viruses and defective virus-like particles, we chose to use the term VLP as a generic name for both fully functional and defective viruses in the rest of the results section.

### sEVs and VLPs can be separated through a velocity gradient

Because the protocol of separation shown in Fig.1A resulted in a mix of EVs and VLPs in the 10k and 200k, we next tried to separate VLPs from the other EVs. We first used asymmetric flow field-flow fractionation (AF4), which had been previously successfully implemented to separate exomeres (also called ENPs) and two types of EVs, respectively of 60-80 and 90-120 nm of diameter (Zhang et al., 2018). Using settings shown in Suppl. Fig.2A, we managed to separate from 200k pellets of EO771 small protein-rich structures (Suppl. Fig.2B: P2) from light-scattering EVs, among which we recovered three populations of 35-80 nm (P3), 80-180 nm (P4) and 180-220 nm (P5) diameters (Suppl. Fig.2B-C). However, this method did not separate VLPs from small EVs of the same size, since both were identified by cryo-EM in the P4 and the P5 EV-containing AF4 fractions of the 200k pellets (Suppl. Fig.2D).

**Figure 2:**
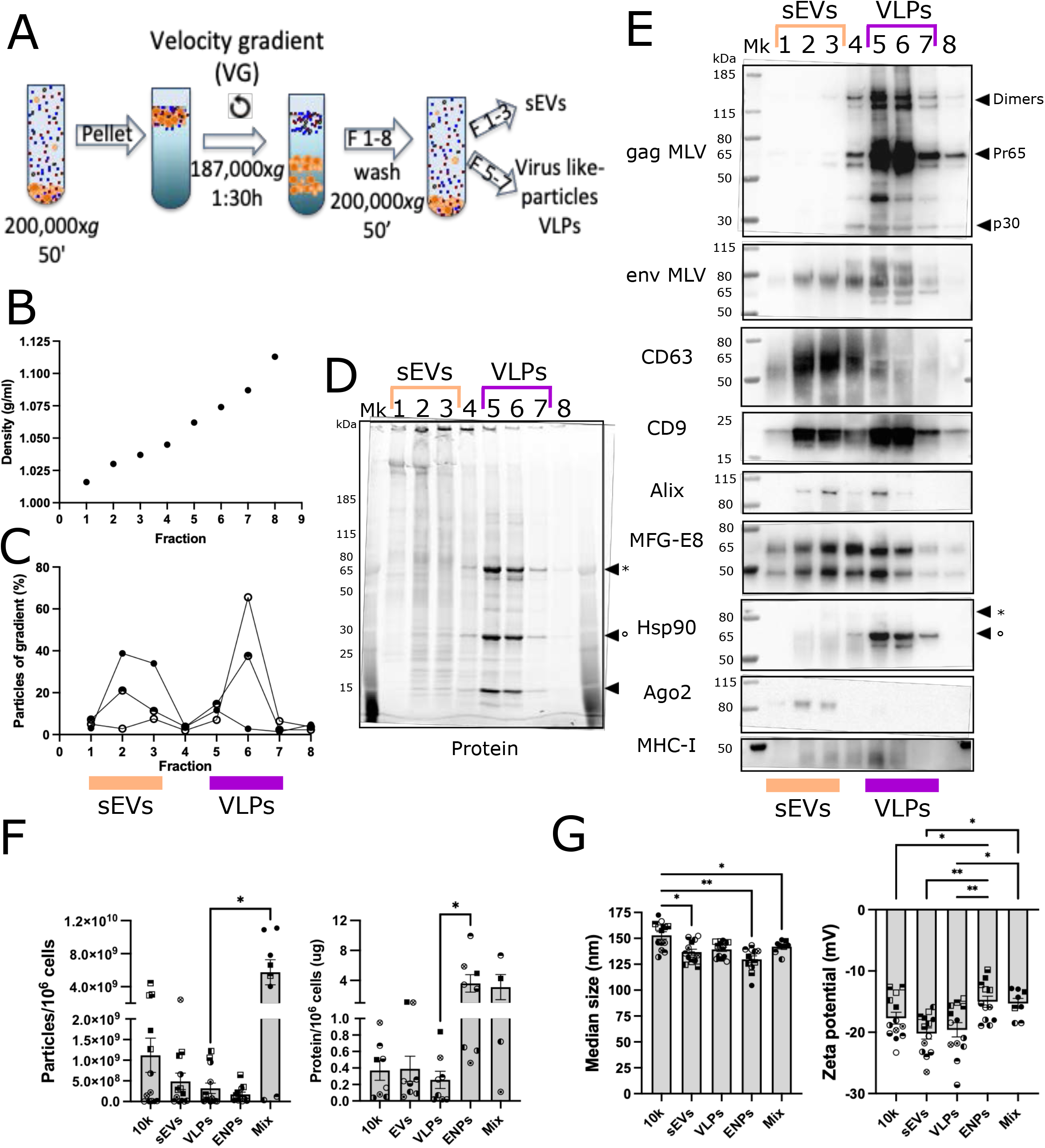
Separation of sEVs from VLPs. (A) Scheme of the protocol of separation of sEVs and VLPs starting from the 200k pellet. (B) Density of fractions of 2ml (1-8) of the velocity gradient. (C) Percentage of number of particles in each fraction over the total measured for all the gradient. n=3. (D) Protein stain-free image of a representative gel of fractions 1-8 from the gradient. * ° = prominent bands in fractions 5-7 of the same size as p65 and p30 gag bands in 2E. (E) Representative western blot stained with antibodies against MLV, EV and ENP markers. * ° in Hsp90 = position of full-length and short bands as observed in the 10k and 200k of Fig.1B. (F) Particle number (left) and protein amount (right) in each subtype of particle (10k, sEVs, VLPs, ENPs and Mix) secreted by 10^6^ EO771 cells. n=13. (G) Median size (left) and zeta potential (right) of each subtype of particle. n=13. Statistical analysis. *: p< 0.05; **: p<0.01.

We then refined the ultracentrifugation protocol by adding an additional step of velocity gradient (Fig.2A). We loaded the 10k (Suppl. Fig.3) or 200k (Fig.2) from EO771 cells on top of an iodixanol density gradient and centrifuged for a short time (1h30), adapting a protocol previously performed to separate HIV-1 virus from other EVs (Cantin et al., 2008; Liao et al., 2019; Martin-Jaular et al., 2021). After measuring their density (Fig.2B), we washed eight fractions of 2 ml each (1 to 8, top-to-bottom) by ultracentrifugation, resuspended them in PBS and processed for NTA measurements (Fig.2C) and western blot analysis (Fig.2D-E). Two peaks of particles were observed from the 200k: in fractions 1 to 3 and in fractions 5 to 7 (Fig.2C), while from the 10k, a single peak of particles was observed in the denser part, fractions 5 to 7 (Supp. Fig.3A). The total protein stain of the 200k fractions showed distinct patterns of proteins in fractions 1 to 3 and 5 to 7 (Fig.2D, Suppl. Fig.3B). Remarkably, 3 prominent bands were detected in the denser fractions (5-7) but absent in the low-density fractions (1-3). We then probed the western blot with an antibody against gag of MLV (Fig.2E, Suppl; Fig.3B), revealing several bands in fractions 5 to 7, 2 of which likely corresponded to the prominent proteins (around 65kDa and 30kDa, * and ° in Fig.2D). The env MLV protein was also enriched in fractions 5-7, displaying different size variants, but it was also found in fractions 1-3 (Fig.2E, Suppl. Fig.3B). The EV markers were distributed along the 8 fractions with different abundances depending on the marker (Fig.2E). While CD9, MFGE8 and Alix were equally present in fractions 1-3 and 5-7, CD63 and Ago2 were mostly in fractions 1-3, whereas MHC-I and the short band recognized by anti-Hsp90 (° in Fig.2E, Hsp90 panel) were most prominently detected in fractions 5-7. These results show that fractions 1 to 3 from the velocity gradient-based separation of 200k contained mainly endogenous and virus-modified sEVs, while fractions 5 to 7 contained VLPs. Fraction 4 consisted in a mixture of both and fraction 8 was mostly empty. By contrast, the velocity gradient was not effective to separate the EVs from the VLPs in the 10k pellets, since all the particles were recovered in fractions 5-7 (suppl Fig.3A, B). Velocity gradient applied to 200k from Pfa1 cells showed absence of the prominent bands and of gag-labeled proteins in fractions 5-8, confirming that these cells do not release VLPs (Suppl. Fig.3C).

Thus, using a pipeline of combined protocols described in Fig.1A and 2A (Supp. Fig.3D), we separated 3 different types of particles or vesicles: sEVs (200k velocity gradient fractions 1-3), VLPs (200k velocity gradient fractions 5-7), and ENPs (200,000xg ON pellet of the supernatant of 200,000xg 50 min, Fig.1A, Supp. Fig.3D). Additionally, we isolated two different mixtures of EVs and/or ENPs: the 10k pellet (10k) containing large/medium and small EVs plus VLPs, and a mixture of all the remaining particles pelleted by ultracentrifugation at 200,000x*g* ON of the supernatant of the 10,000x*g* centrifugation (Supp. Fig.3D). This new sample, called “Mix”, was included in the following experiments, as representative of an heterogenous mixture of particles physiologically secreted by tumor cells. We systematically recovered these 5 sample types (10k, sEVs, VLPs, ENPs, Mix) from each batch of EO771-conditioned medium, in several independent experiments. We quantified the number of particles by nanoparticle tracking analysis (NTA) and the amount of proteins recovered in each sample (Fig.2F). The Mix fraction was the most enriched in particles, while ENPs were the least enriched. By contrast, the amount of proteins was the highest in the ENPs and in the Mix. Analyzed by NTA (Fig.2G), the 10k had a significantly larger median size than sEVs and ENPs, but not significantly different from that of VLPs. The zetapotential of ENPs was significantly less negative than that of all EVs types (Fig.2G).

### Quantitative proteomic analysis of the different particle subtypes

We next performed a quantitative proteomic analysis by label-free mass spectrometry on 10k, sEVs, VLPs, ENPs and Mix samples from EO771, adjusted to the same amount of proteins. Proteins were identified combining the databases of mouse and murine virus proteins as in Fig.1F. 6127 proteins in total, including 53 viral proteins, were identified by at least 3 peptides among 5 replicates in at least one type of sample (Suppl. Table 2: VennDiagram). Venn diagram analysis showed that most of the identified proteins (3800) were shared between all groups (Fig.3A, with Mix sample excluded from the Venn Diagram analysis for better visualization of the single fraction-specific components). Nevertheless, 303 proteins were found only in the 10k, 130 in the sEVs, 63 in the VLPs and 231 in the ENPs (Suppl. Table 2: Venn Diagram). Principal Component analysis (PCA) of the label-free quantification (LFQ) results (Suppl. Table 2: LFQ), showed clustering of the 5 replicates of each fraction (Fig.3B, and heatmap of protein abundance: Suppl. Fig.4A), confirming the reproducibility of our protocol. ENPs and the Mix were tightly close, indicating the prevalence of ENPs in the latter. Both were far from the membrane-containing 10k, EVs and VLPs, although to a lesser extent for the Mix, in agreement with the heterogeneous composition of this fraction which contains sEVs, VLPs and ENPs. Because of this mixed composition, the Mix was excluded from further analyses, since we aimed at identifying specific protein signatures of EV subtypes.

**Figure 3:**
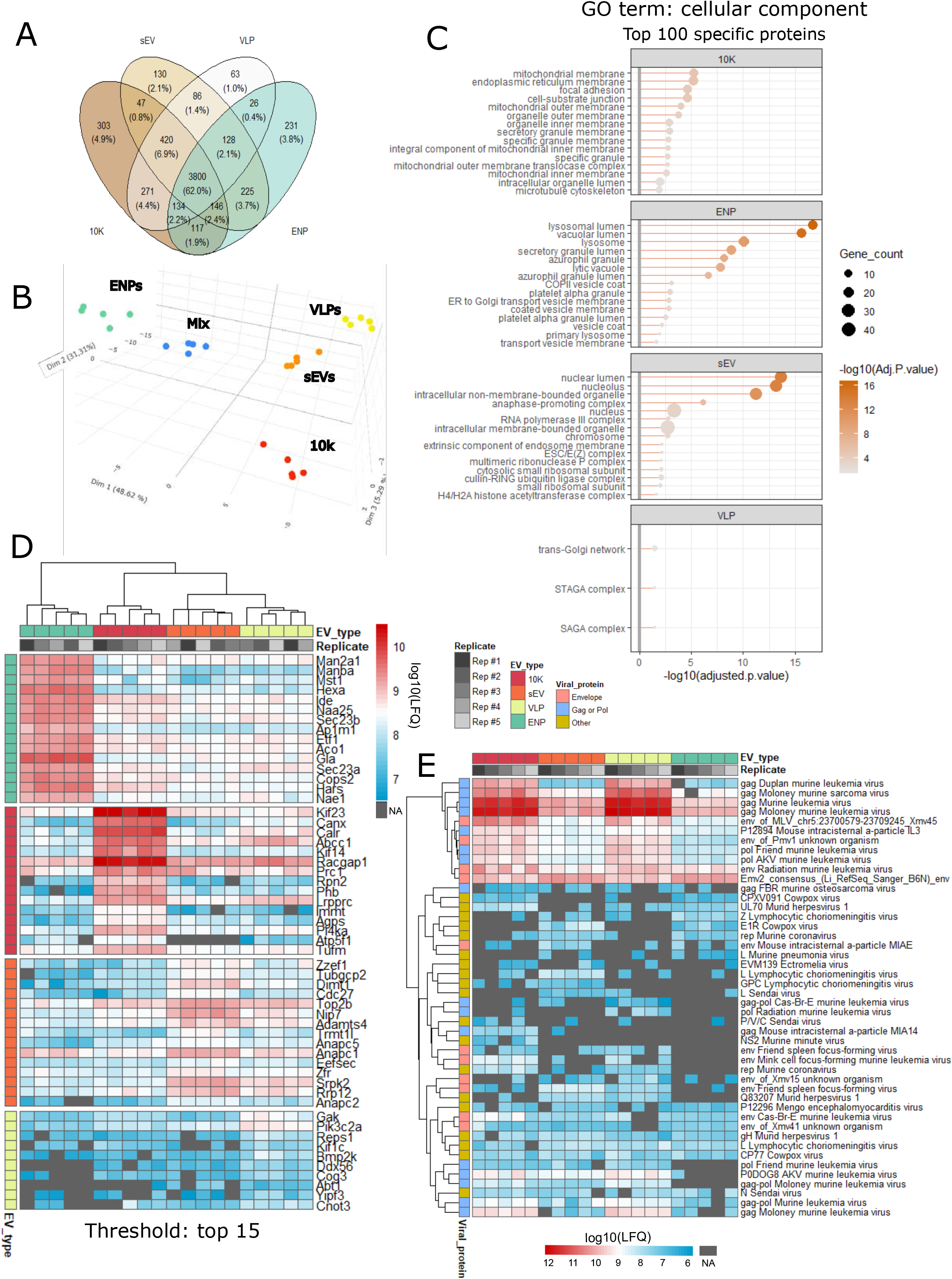
Proteomic characterization of subtypes of particles. (A) Venn diagram of all (mouse + viral) proteins identified with at least 3 peptides among 5 replicates in at least one sample type among the 10k, sEVs, VLPs and ENPs. All proteins are listed in Suppl. Table 2, tab “VennDiagram”. (B) 3D view of the first 3 components of the principal component analysis (PCA) of all samples based on their protein abundance (LFQ). Samples are colored by sub-types: 10k = red; sEVs = orange, VLPs = yellow, ENPs = green, Mix = blue. (C) GO term-enrichment analysis for cellular components in the top-100 most specific proteins (from Suppl. Table 2, SSPA): the only 3 significant GO-terms (for VLPs) or the 15 most significant GO-terms (for 10k, sEVs and ENPs) are shown. GO-terms are ordered by adjusted.p-value. (D) Heatmap of protein abundance of the 15 most specific proteins (or the only 10 specific proteins for VLPs) of each group (from SSPA analysis, Suppl. Table 2). Clustering is shown to confirm that specific proteins are not identified due to outlier replicates. NA= not detected/absent. (E) Heatmap of protein abundance of the 46 quantified viral proteins in the different fractions. All gag-pol sequences are primarily present in the 10k and VLPs, whereas envelope proteins (env) are more equally distributed between the fractions. NA= not detected/absent.

A Gene Ontology (GO)-term enrichment analysis was performed, first on the top-100 most abundant proteins of each group (i.e. including the shared abundant proteins) (Suppl. Fig.4B). The most significantly enriched “cellular component” GO-terms found in all samples were focal adhesion and cell-substrate junction, suggesting that areas of cell interaction with each other and with their substrate were major sources of EVs/ENPs. Other enriched GO-terms included ribosome-and nucleus-related terms, which were especially significantly enriched in sEVs and in ENPs, while terms corresponding to lumen of various internal compartments (including ficolin-1-containing compartments) were more enriched in 10K and VLPs.

We next used the state-specific protein analysis (SSPA) feature of the myProMS software (as described in Materials and Methods) to identify proteins specific to or significantly enriched in one or a group of samples (Suppl. Table 2: tab “SSPA”). GO-term enrichment analysis on the top-100 most enriched specific proteins of each sample highlighted membranes of mitochondria, ER and secretory granules in the 10k, lumen of secretory granules and lysosomes as well as membrane of secretory and coated vesicles in ENPs, and nuclear components as well as both lumen and membrane components of organelles in sEVs. Given the low number of specific proteins in VLPs (see Fig.3A, 3D), only 3 significantly enriched GO-terms were identified: trans-golgi network, and two complexes involved in regulation of transcription and histone modification (the STAGA and the SAGA complexes) (Fig.3C). Abundance of the 15 most specifically enriched proteins of each sample type (only 10 for VLPs) is shown in Fig.3D.

Finally, when we focused the analysis on the identified viral proteins (Suppl. Table 2), the majority of peptides matched with MLV proteins as identified by a few proteo-specific peptides: AKV, Duplan, but also endogenous MLV (MLV-chr5). Other viruses identified by more than one specific peptide included intracisternal a-particle, and, in lower amounts, Cas-Br-E MLV, Xmv41, lymphocytic choriomeningitis virus, murid herpes virus, mengo encephalomyocardiatis virus and murine coronavirus. The SSPA (Suppl. Table 2, Fig.3E) showed that gag-pol and pol proteins were generally enriched in both the 10k and VLPs as compared to sEVs and ENPs. The envelope proteins, by contrast, were variably distributed in all fractions. These distributions, therefore, confirmed the western blot analyses (Fig.1G, 2D, Suppl. Fig.3B). A few other viral proteins were detected, although at low levels, in sEVs and ENPs: the RNA polymerase L of the sendai and the lymphocytic choriomeningitis virus, or the replicase of coronavirus, suggesting that elements of other viruses can end up in sEVs.

### Tumor derived EVs and particles induce distinct phenotypic changes in DC

Given the different protein composition of each type of particles, we hypothesized that they may induce different effects on target cells. Due to our primary interest for the role of tumor-derived EVs in establishment of antigen-specific immune responses, we chose canonical antigen presenting cells, dendritic cells (DCs), as targets of tumor-derived EVs for the subsequent functional studies. The same amount of proteins (10 or 20 µg/ml final) of each of the particle fractions produced by EO771 and characterized above were added for 16 h to MutuDC, a model cDC1 cell line from C57BL/6 (Fuertes Marraco et al., 2012). Then, we measured MutuDC cell viability and surface expression of maturation markers by flow cytometry, and we quantified cytokine release in the supernatant by cytokine-bead array (CBA) (Fig.4A).

**Figure 4:**
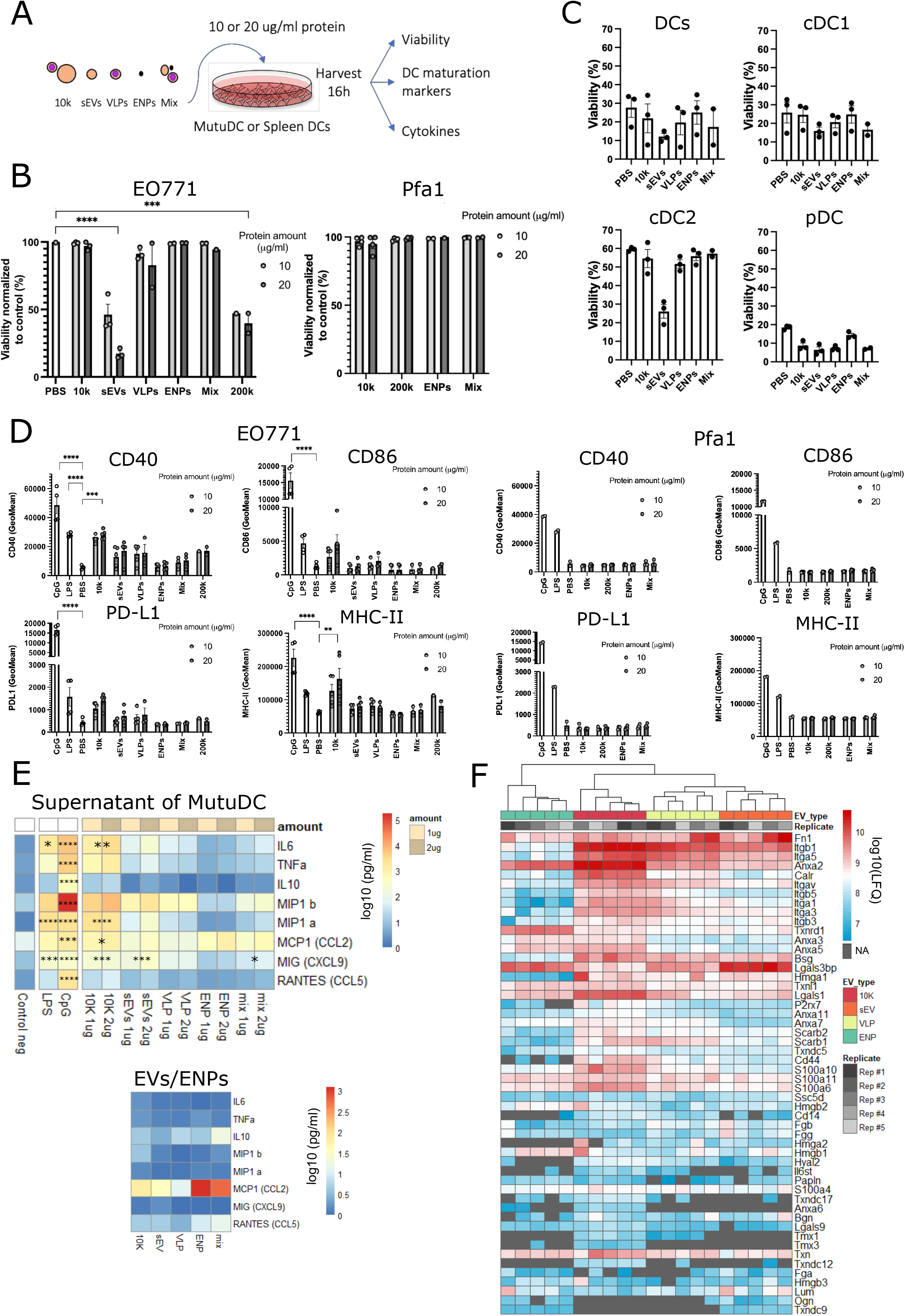
Phenotypic changes induced by the different subtypes of particles from EO771 on MutuDC and spleen DC. (A) Scheme of the in vitro experiment and the parameters assessed. (B) Viability of MutuDC cells after 16h of exposure to 10 or 20 µg/ml of the different particles coming from EO771 tumor cells (left) or the non-tumor cell line, Pfa1 fibroblasts (right), measured by flow cytometry with eFluor 780 fixable viability dye, normalized to the control non-treated condition. n=3. (C) Viability of isolated spleen DCs from C57BL/6 mice after 16 h of exposure to 20 µg/ml of the different particles coming from EO771tumor cells. n=3. Results are shown for total DCs (CD11c+), cDC1 (XCR1+), cDC2 (CD172a+), pDC (B220+) as indicated above each graph. (D) Expression of maturation markers on viable MutuDC cells after 16 h of exposure to 10 or 20 µg/ml of the different particles coming from EO771 cells (left) and Pfa1 cells (right), measured by flow cytometry (GeoMean). LPS and CpG treatments were used as positive controls of maturation. n=5. (E) Quantification of cytokines secreted by DCs in the supernatant of MutuDC exposed to the subtype of particles (top) and in the particles themselves (bottom), presented as heatmap (log10 scale). “Amount 1 µg/2 µg” refers to the amount of EV/ENP used to treat MutuDC (in 100 µl final volume). n=3. Statistical analyses were performed using mixed-effects model with Dunnett’s multiple comparison to the PBS, the mean of both concentrations was used for the comparisons, with different concentrations as repeated measures (B, D and E). *: p<0.05, **: p< 0.01, ***: p< 0.001 and ****: p< 0.0001. (F) Heatmap of protein abundance of proteins qualifying as DAMPs and PAMPs as quantified by LFQ in our proteomic analysis of the different types of particles (Suppl. Table 2). NA= not detected/absent.

Remarkably, purified sEVs induced the death of MutuDC (Fig.4B, left, flow cytometry gating strategy shown in Suppl. Fig.5A). This toxicity was not due to insufficient washing of the iodixanol from the fractions 1-3, since it was not observed with the higher density iodixanol gradient fraction VLPs, while the 200k, containing sEVs but devoid of any iodixanol, also induced death (Fig.4B, left). We observed a very limited cytotoxic effect induced by either the 10k, ENPs or Mix, and a slight toxicity (but still much weaker than that of sEVs) induced by VLPs. This cytotoxicity was not observed at all when using EVs and particles isolated from the non-tumor cell line Pfa1 (Fig.4B, right).

To confirm these results in a more physiological context, the very same treatments were applied to primary DCs isolated from the spleen of a C57BL/6 mouse (Fig.4C, gating strategy in Suppl. Fig.5B). Basal viability of isolated DCs after 16 h of culture was low (PBS condition in Fig.4C): 20-30% for the global DC population, with a better survival of cDC2 (60% viability). Treatment by sEVs (but not by any other samples) further decreased the viability to 10% for the global population and to 20% for cDC2. The low survival rate of Pdc (20%) was decreased to 10% by treatment with any vesicle-containing samples (10k, VLPs, sEVs, Mix), but not by treatment with ENPs.

We then asked how EV/ENP subtypes changed the maturation profile of MutuDCs (Fig.4D, Suppl. Fig.5A). CpG and LPS were used as internal control of DCs’ ability to respond to maturation-inducing signals. Treatment by the 10k pellets significantly increased expression of CD40 and MHC-II and, to a lower (and not significant) extent, of CD86 and PD-L1 (Fig.4D, left). Treatment by sEVs, VLPs and 200k only slightly increased expression of CD40 and MHC-II, whereas ENPs did not induce increase of any of these markers (Fig.4D, left). MutuDC did not overexpress any of the analyzed co-stimulatory molecules when exposed to the Pfa1 EVs or particles (Fig.4D, right).

A panel of 17 cytokines reported to be secreted by DCs were next quantified in the conditioned medium of MutuDC. Eight of these cytokines were detected in at least one of our samples (Fig.4E). Three of them were also detected in the EVs and ENPs that had been used for treatment of MutuDC (bottom panel Fig.4E): in particular, MCP1 was abundant in ENPs and also present in the other particles, while RANTES and IL10 were detected in all types of EVs and in ENPs but at very low level. Therefore, MCP1 detected in the supernatant of all EV-treated MutuDC was thus possibly coming from the EVs rather than from the cells, and will not be discussed further. The 10k pellet was the most efficient to induce cytokine secretion by MutuDC, which released the 7 cytokines at a higher level than control (untreated) MutuDCs: 3 for which the difference with control treatment was statistically significant (IL6, MIP1a, MIG), 2 which were not statistically different from control but still expressed at a much higher level (TNFa, MIP1b), and 2 expressed at low level (IL10, RANTES). The second most efficient treatment was sEVs, which induced statistically significant increased secretion of MIG, and non-statistically significant (but still higher than control) levels of IL6, TNFa, MIP1b, MIP1a, RANTES. However, given the high rate of cell death observed in these cultures, some of these cytokines could have been induced by detection of cell death rather than by direct signaling induced by sEVs. VLPs induced some IL6, MIP1b and MCP1, and even smaller amounts of TNFa and MIP1a. The only cytokine detected in ENP-treated DCs was MCP1, probably coming from ENPs themselves.

In conclusion, the 10k was the most inflammatory treatment, triggering the maturation of MutuDC and the secretion of several pro-inflammatory cytokines and chemokines, whereas ENPs did not have any effect on MutuDCs, VLPs induced low levels of DC maturation, and sEVs induced some maturation and cytokine secretion, but also significant cell death. Consistent with this observation, our quantitative proteomic data showed that proteins qualifying as DAMPs and PAMPs (according to a list defined as such in (Hoshino et al., 2020)) were most abundant in the 10k pellet (Fig.4F).

### Membrane-containing particles 10k, sEVs and VLPs, but not ENPs can transfer protein effectively to DCs

A proposed mode of action of EVs is to deliver their content into target cells following their uptake. We thus wanted to compare the capacities of uptake of each of the EVs and particle subtypes. We chose the fluorescent protein, mCherry as cargo of EVs/ENPs, as it is amenable to visualization and quantification by fluorescence microscopy, flow cytometry and spectroscopy. To induce its loading into EVs/ENPs, we introduced a myristoylation and palmitoylation sequence upstream of mCherry, to generate the myr/palm-mCherry (Fig.5A) (Mathieu et al., 2021; Valenzuela and Perez, 2020). In the resulting protein, post-translational addition of the acyl chains that insert into the cytosolic side of the lipid bilayer should lead to association of the fluorescent protein with internal membranes or lipidic structures (McCabe and Berthiaume, 1999, 2001) (Fig.5A), resulting in the targeting of this membrane-bound mCherry inside the released EVs and possibly in/on ENPs.

**Figure 5:**
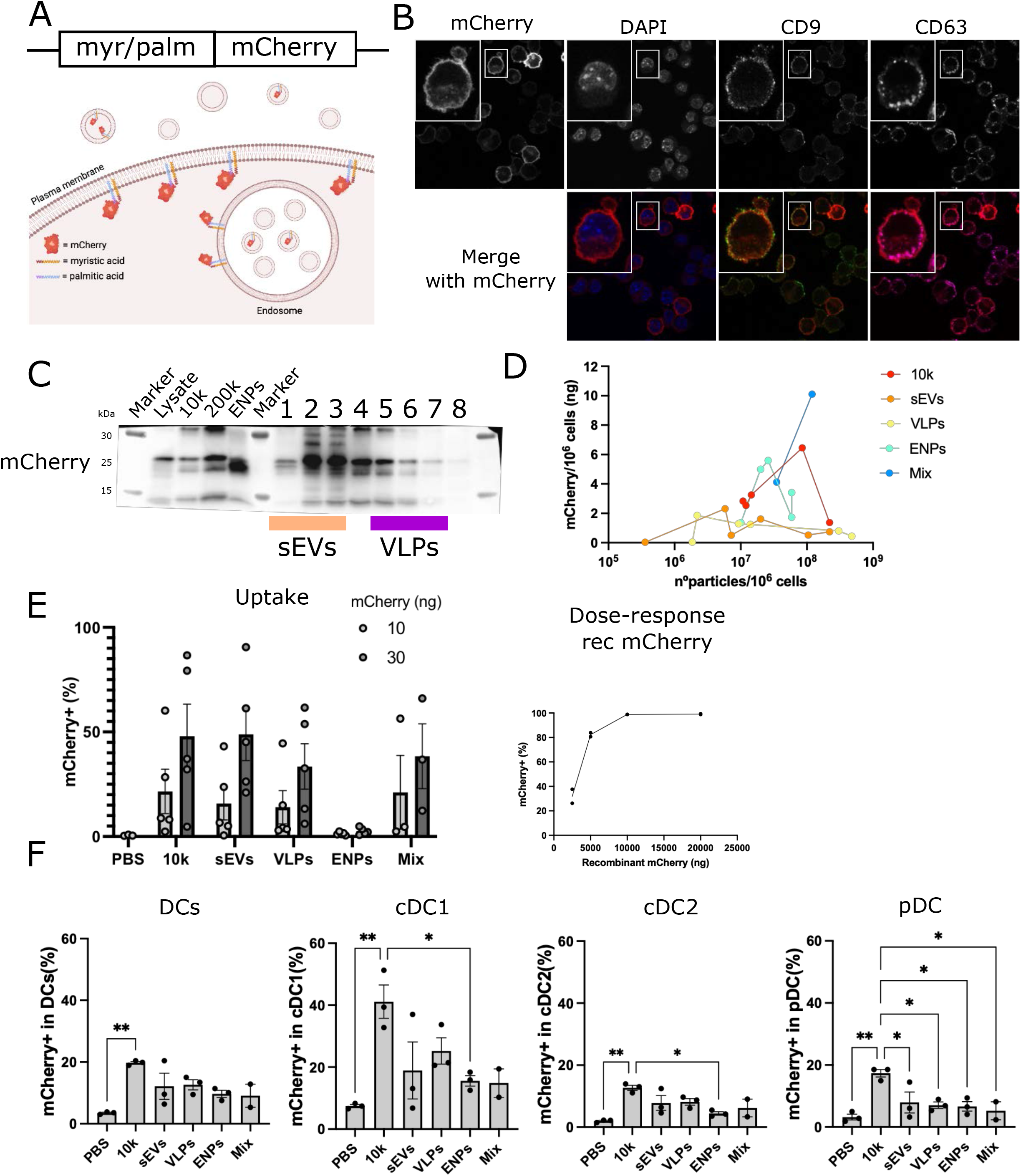
Protein transfer capacity of the different subtypes of particles. (A) Scheme of the construct containing the myristoylation and palmitoylation sequences fused to mCherry, which was used to create the EO771 myr/palm-mCherry stable cells (top) and scheme of expected intracellular and intravesicular distribution of myr/palm-mCherry (bottom, created by Biorender.com). (B) Confocal microscopy of EO771-myr/palm-mCherry cells showing DAPI in blue and mCherry in red, CD9 in green and CD63 in magenta (overlay and close-ups). (C) Western blot loaded with EVs/ENPs from 20×10^6^ cells hybridized with anti-mCherry, showing its presence in the different subtypes of particles (same gels as in Fig.1C,G and Fig.2D,E). (D) mCherry fluorescence quantification (excitation 585/emission 625) measured with a spectrophotometer for each preparation of particle and expressed in ng, as calculated from a standard curve made with known concentration of recombinant mCherry. Results are presented as a function of the number of particles quantified by NTA in the same preparation. n=6. (E) Uptake by MutuDC, quantified by flow cytometry (expressed as % of mCherry+ cells), of 10 or 30 ng of mCherry from each subtype of particle (left, n=5), or a dose-response of recombinant mCherry (middle, n=2). (F) Uptake by spleen DCs of 30 ng of mCherry from each subtype of particle. Results are presented for total CD11c+ DCs, and subtypes of DCs: cDC1, cDC2, and pDCs (defined as in Fig.4 and suppl. Fig.5). One way ANOVA with Tukey’s multiple comparison test was performed, n=3. *: p<0.05 and **: p<0.01.

We transfected EO771 with a myr/palm-mCherry-expressing plasmid and we sorted mCherry^+^ cells by flow cytometry to generate a stable cell line. We checked the cellular localization of the mCherry in the cells by confocal microscopy (Fig.5B). The mCherry fluorescence was found mainly at the plasma membrane of the cells but also internally, colocalizing partially with both CD9 and CD63.

To determine the targeting of the mCherry into the particle subtypes, the same western blot from Fig.1 and Fig.2 (which contained EV/ENP samples from myr/palm-mCherry-EO771 EVs) was revealed with an antibody against mCherry (Fig.5C): two bands around 25kDa were detected in all particles as well as in all gradient fractions. mCherry was also detected in the ENPs, mainly as the lower band form. mCherry fluorescence was measured in each sample and compared to a standard curve of recombinant mCherry to quantify the absolute amount of mCherry (Fig.5D). Since mCherry amount was not linearly correlated with the number of detected particles (Fig.5D), we chose for each type of sample to use the same amount of mCherry as quantified by fluorescence, rather than the same particle number or protein amount, to feed target cells and measure efficacy of uptake.

When exposed for 16h to 10 or 30 ng of EV/ENP-associated mCherry or recombinant mCherry (rec mCherry), MutuDCs were able to incorporate the fluorescence from the 10k, sEVs, VLPs and Mix similarly and in a dose dependent manner, but not from the ENPs (Fig.5E, left). Addition of a 100-fold excess of recombinant protein (2500 ng, Fig.5E, middle) was required to achieve the same degree of uptake observed with 30 ng of EV-associated mCherry.

We next performed the uptake assay with primary spleen DCs, which were exposed to 30 ng of EV/ENP-associated mCherry (Fig.5F). In all DC subtypes, the 10k was significantly uptaken, as compared to the PBS control and to ENPs, whereas the VLPs, sEVs, and Mix showed some level of uptake, but not significantly different from the PBS control. cDC1, the DC subtype specialized in cross-presentation, were the cells displaying the highest uptake capacity, mainly of the 10k, but also (although not significantly) of other EVs and ENPs. Our results demonstrate that the mixed EV-containing 10k is successfully captured and recognized by DCs, with a better efficiency compared to ENPs or free soluble protein.

### The 10k is the scaffold that leads to the highest cross-presentation

Finally, to directly address the capacity of EVs/ENPs to induce presentation of antigens by target DCs, we used OVA as a model antigen, fused to the same myr/palm sequence used with mCherry, to allow its targeting to EVs/ENPs, and a myc tag for detection (Fig.6A). In EO771 cells stably expressing the resulting protein, OVA was detected as bands of different sizes in the secreted EVs and particles, and most prominently in the 10k (Fig.6B). The amount of OVA was quantified in each sample as compared to known concentrations of recombinant OVA loaded on the same western blot, and rapported to either the number of secreting cells (Fig.6C, top), or to the number of particles (Fig.6C, bottom). Like for mCherry uptake, OVA concentration was used to normalize the amount of each fraction used for the cross-presentation experiments.

**Figure 6:**
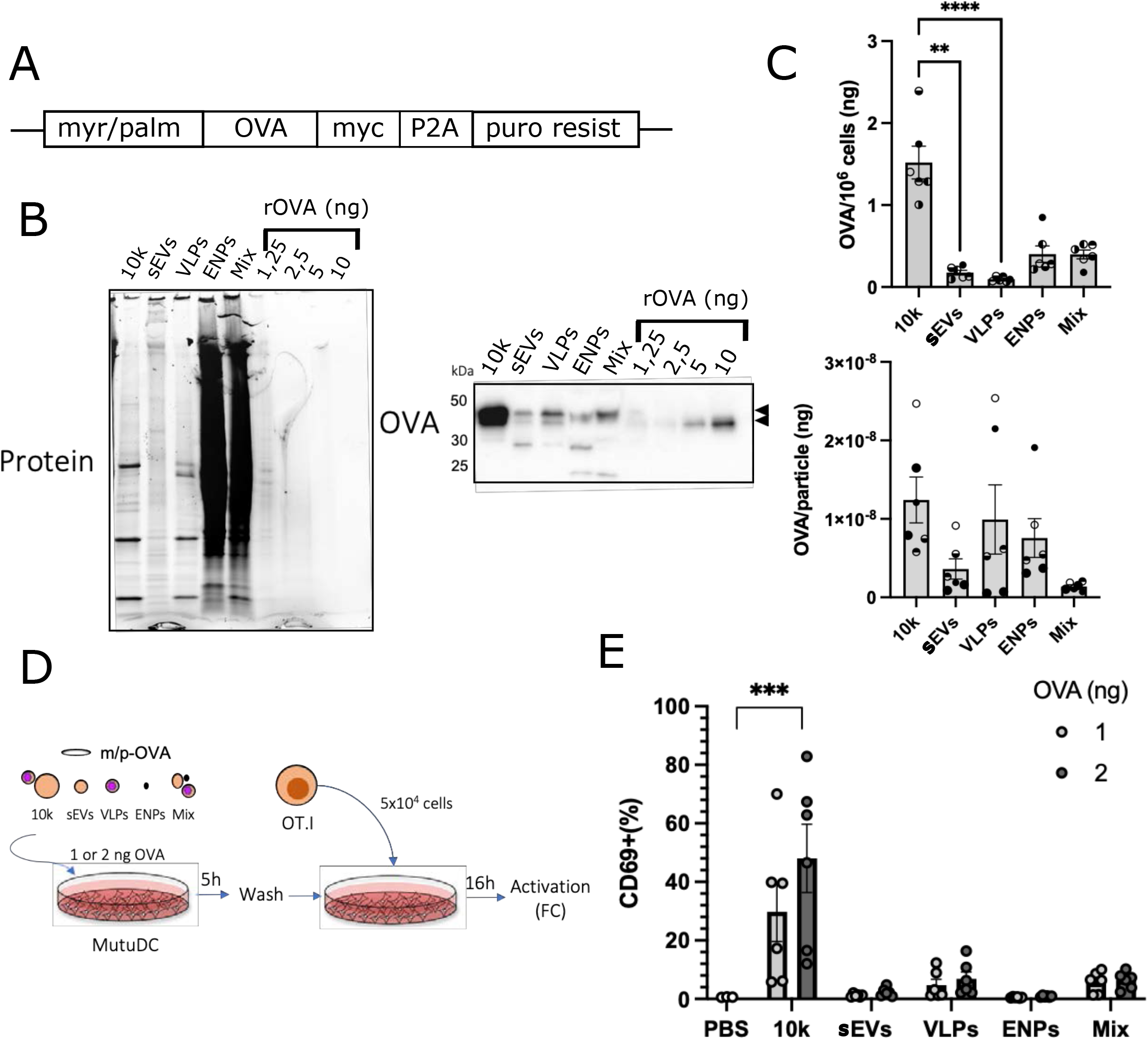
Cross-presentation of OVA antigen carried by the subtypes of particles. (A) Scheme of the construct containing the myristoylation and palmitoylation sequences fused to OVA, myc tag, P2A cleavage site and puromycin resistance gene for selection, which was used to create the EO771 myr/palm-OVA stable cells. (B) Analysis of EVs/ENPs from 20×10^6^ EO771-m/p-OVA by western Blot. Four known amounts of recombinant OVA were loaded on the same gel to allow quantification of OVA. Blots were revealed with a polyclonal antibody against OVA. One representative blot out of 6. (C) Quantification of OVA in the different EVs/ENPs, was done on the western blot images as compared to the OVA dose-response curve. Graphs show ng OVA/10^6^ cells (top) or ng OVA/particles (bottom). Friedman test with Dunn’s multiple comparison was performed. n=6. (D) Scheme of the protocol of cross-presentation of EV/ENP-OVA cargo by MutuDC to OT1 T cells. (E) Activation of OT1 T cells exposed for 18h to MutuDC that had been fed with OVA (1 or 2 ng) present in the various types of EVs/ENPs. T cell activation was quantified as percentage of CD69^+^ cells by flow cytometry. Mixed-effects model was performed with Dunnett’s multiple comparison to the PBS, the mean of both concentrations was used for the comparisons, with different concentrations as repeated measures, n=6. *: p<0.05, **: p< 0.01, ***: p< 0.001 and ****: p< 0.0001.

MutuDC cells were exposed to 1 or 2 ng of OVA associated to the different carriers for 5 h, then washed and co-cultured with OT1 cells, expressing a TCR that recognizes specifically the SIINFEKL peptide of OVA presented on MHC-I H2-Kb molecules (Fig.6D). After 18h, activation of OT1 T cells was measured by flow cytometry, quantifying expression of the early activation marker CD69 (Fig.6E). In this assay, the most efficient (and only statistically significant) activation of T cells was observed in the presence of MutuDC exposed to 10k. MutuDC exposed to VLPs or to Mix induced a very low, but clearly above background (although not significantly different from PBS control), level of OT1 activation, while MutuDC exposed to sEVs and ENPs did not activate at all the T cells.

## DISCUSSION

Tumor-derived EVs have been proposed as a source of tumor antigen and as immune regulators in the tumor microenvironment. However, the bibliography is contradictory, attributing to tumor EVs both immuno-activating and immuno-suppressive roles (Greening et al., 2015; Robbins & Morelli, 2014; Théry et al., 2009). These discrepancies could be explained by differences in the tumor cells used as source of EVs (cell type, cancer stage, culture conditions), but also by the isolation methods, which could result in various mixtures of the EVs and ENPs that a single tumor cell secretes.

In this work, we show that the ENPs and EV subtypes from a mouse mammary adenocarcinoma cell line, EO771, have distinct and even opposite effects on target DCs in vitro. Most strikingly, we evidenced a novel, and thus far ignored, subtype of small EVs: VLPs and infectious retroviral particles, which represent a major component of mouse tumor-derived EV preparations.

By designing a robust protocol to separate the VLPs and retroviruses from other small EVs and comparing side-by-side their effect on target antigen-presenting cells (DCs), we unexpectedly observed that virus-free sEVs induced death of the DCs, whereas VLPs did not display this toxic activity. Both sEVs and VLPs were similarly captured by MutuDCs, where they induced some surface expression of maturation markers and low levels of cytokine secretion. However, despite sEVs led to a slightly higher level of cytokine secretion than VLPs, they did not induce cross-presentation of a model antigen they had been engineered to carry, while VLPs did allow some cross-presentation. Therefore, some of the literature describing immune effects of murine tumor-“exosomes” should be reconsidered, as describing effects of a combination of toxic sEVs (possibly coming from both MVBs and the plasma membrane) and antigen-carrying VLPs and retroviruses.

The VLPs, found here in three tumor cell lines (EO771, 4T1, MutuDCs), come from endogenous retroviruses (ERVs) with restored infectivity. The presence of endogenous retroviral proteins has been reported in mouse tumor cell lines (Leong et al., 1988; Shepherd et al., 2003) and in EV preparations of long-term cultures of a mouse dendritic cell line (Thery et al., 2001; Thery et al., 1999). In fact, ERVs are expressed in mice at low levels, but they are non-infectious (Stocking and Kozak, 2008). However, a particular ERV locus of C57BL/6 mice, *Emv2*, encoding for an endogenous ecotropic replication-defective murine leukemia virus (eMLV) carrying a single inactivating mutation in the reverse transcriptase, can by *trans*-complementation of viral proteins and eventually recombination with other endogenous MLV proviruses, become infectious and subsequently expressed at high levels (King et al., 1988). In the organism of C57BL/6 mice, the acquisition of infectious potential of this gene is stopped by the immune system in an antibody-dependent manner (Young et al., 2012), but in *in vitro*-propagated cell lines this control process does not take place. In fact, analysis of a panel of tumor cell lines derived from C57BL/6 mice showed that most of them expressed infectious eMLV at high levels (Ottina et al., 2018). Since we also observed VLPs produced by a mouse cell line of Balb/C origin (4T1), it is likely that EVs from most mouse tumor cell lines used to study cancer development and therapeutic approaches contain VLPs and/or infectious retroviral particles. It may not be the case for most human tumor-derived cell lines, except those established from retrovirus-induced cancers, since endogenous retroviruses are mostly silent in the human genome (Kassiotis and Stoye, 2017). However, human endogenous retrovirus (HERV)-derived RNA and DNA have been previously detected in EVs from human gliomas and medulloblastomas (Balaj et al., 2011), and there are several reports of retroviral particle production by human placenta and cancer cells (Contreras-Galindo et al., 2008; Keydar et al., 1984; Lower et al., 1993; Nelson et al., 1978), thus calling for a more systematic assessment of the presence of VLPs and infectious retroviruses in EVs released by human tumor cell lines.

Regarding the immunogenicity, our work shows that the VLP-containing EV preparations, 10k and VLPs, are the most immunogenic for DCs, promoting their maturation, cytokine and chemokine secretion, cross-presentation and activation of T cells. This may be due to a viral-dependent mechanism, such as viral RNA-or DNA-mediated activation of pattern recognition receptors (PRRs). However, we did not observe release of IFN type I cytokines by DCs, suggesting that another pathway may be activated by these VLPs. Further mechanistic studies would be required to determine the nature of the activation pathways induced by VLPs in DCs.

Importantly, the 10k and 200k of a non-tumor cell line, Pfa1, did not induce maturation of DCs, as opposed to the tumor-derived 10k, VLPs and sEVs, nor cell death, as opposed to tumor-derived sEVs. This suggests that tumor-derived EVs have some immune functions that are tumor-specific, and that DCs are able to differentiate tumor-from non-tumor EVs. One possible reason for this difference is the expression of complete and infectious eMLV. Indeed, an env protein was detected in Pfa1 EVs, but not gag, and the dense fractions of the velocity gradient were empty, suggesting that there were no VLPs formed by this cell line (Suppl. Fig.1,3). For the specific toxic effect of sEVs, however, other gag-independent and tumor-specific cargoes must be involved. For instance, DAMPs, that might be produced by the genomic deregulation characteristic of tumor cells, may be such cargoes. The list of proteins specific or strongly enriched in one or the other subtype of EVs, provided by our work (Fig.3-4, Suppl. Table 2), will be a very useful source to explore for candidate proteins responsible for these EV subtype-specific activities. However, other types of molecules, such as nucleic acids or small metabolites, could also be involved in these effects.

Overall, the 10k induced much stronger responses than the VLPs. The quantitative proteomic analysis of DAMPs reveals the presence and enrichment of different sets of DAMPs in each subtype, but with the most rich and abundant set in the 10k, suggesting that one or several of these proteins are involved in the activation of DCs. This enrichment could be explained by the fact that the 10k includes large EVs that might contain mitochondria or mitochondria-derived vesicles (according to the GO-term enrichment analysis, Fig.3C), or that might be plasma membrane-derived apoptotic bodies (ApoBDs, 500 nm −2 μm) coming from dying cells in the culture. ApoBDs and mitochondria are rich sources of DAMPs, and ApoBD expose phagocytotic signals and, under stress condition, induce immune activation (Caruso and Poon, 2018; Krysko et al., 2012). In addition, the proteomic analysis showed that a few sequences of env proteins were differently enriched in the 10k versus the sEVs or the VLPs, such as env of Mink cell focus-forming virus, which could confer a higher capacity to fuse and thus deliver the EV content into the target cell.

By contrast, the ENPs did not induce any detectable effects on the immune state of the DCs, maturation or cytokine release. This is probably due to their poor efficacy of being captured by MutuDCs, as demonstrated in our uptake assay, as opposed to the capacity of 10k, sEVs and VLPs to be uptaken (Fig.5). Therefore, even though other groups have shown that ENPs or supermeres can transfer to epithelial cells functional enzymes or receptors (Zhang et al., 2019; Zhang et al., 2021b), this activity does not seem efficient or useful on MutuDCs. Similarly, we and others have shown that the ACE2 enzyme, which is also a receptor for the SARS-CoV2 virus, is present on sEVs but also as a soluble form (Cocozza et al., 2020) or in exomeres (Zhang et al., 2021a). However, even though all forms can bind the virus (Zhang et al., 2021a), the EV-associated form is much more efficient to decrease viral infectivity (Cocozza et al., 2020). Therefore, at least for a purpose of immunogenic stimulation and antigen transfer, or for decoy activities, ENPs, like soluble recombinant proteins, are not the most promising tools.

We postulate that the presence of a lipid bilayer on all EV types makes them good targets for efficient phagocytosis. There is plenty of evidence in the literature that a lipid bilayer promotes recognition and uptake, from classic technics of cells transfection through liposomes (Lasic and Papahadjopoulos, 1995) to specific studies of uptake of nanoparticles (Mosquera et al., 2018). In addition, in EVs, the lipid bilayer contains transmembrane receptors and ligands that have counterpart receptors in membranes from target cells, thus facilitating capture. Besides these major factors, some others might be potentiating this effect. For instance, introducing negative charges on soluble OVA antigen was shown to promote its scavenger receptor-dependent uptake by DCs (Shakushiro et al., 2004). Interestingly, we observed that all the EV/ENP subtypes were negatively charged, by measure of their zetapotential (Fig.2). However, the ENPs were the least electronegative and most poorly uptaken. Another feature of all the highly uptaken EV subtypes is the presence of various envelope protein of MLV. This protein can be glycosylated, which makes it recognized by lectin receptors abundantly expressed by both CD8a+ DCs (cDC1) and CD8-DCs (cDC2). In addition, the lectin receptors could also recognize other glycosylated proteins, abundant on EVs (Williams et al., 2018).

The requirements to induce efficient CD8 T cell activation, for instance to initiate antigen-specific anti-tumor immune responses, are that professional APCs (i.e. DCs) capture and present the tumor-derived antigen, together with co-stimulatory molecules on their surface, while secreting cytokines that will allow polarization of the T cells (Alloatti et al., 2016). Expression of the co-stimulatory molecules and the activating cytokines by DCs, i.e. maturation, are driven by their sensing of DAMPs and PAMPs (Gardner and Ruffell, 2016; Kapsenberg, 2003). Our results show that the 10k allows implementation of all these steps in the most efficient manner. The small EVs, by contrast, induce low levels of activation of DCs, and only the VLPs are able to transfer their content while not killing the cells, thus resulting in some extent of cross-presentation, whereas VLP-free sEVs transfer their content but kill DCs. In previous studies where tumor-derived “exosomes” were used to transfer antigen and induce immune responses, the actual structures responsible for this activity were therefore most likely VLPs.

In conclusion, our work has important consequences for translation of EVs to cancer immunotherapy: a mixed preparation of large- and small-size EVs, including VLPs, corresponding to the 10k used here, seems a more efficient source of activity than highly purified small EVs or exosomes. However, this should be tested for the different producing cell sources, and the presence of infectious murine ERV, as potential relevant factor for immune activation, would have to be demonstrated as not inducing adverse effects in humans. Conversely, our observation of the cytotoxicity of small EVs opens a new route of immunotherapy, by blocking the immunosuppressive activity of the cytotoxic tumor sEVs.

## METHODS

### Cells

All cells were kept at 37°C in a humidified atmosphere with 5% CO_2_. The medullary breast adenocarcinoma cell line EO771 isolated as spontaneous tumor from a C57BL/6 mouse (Homburger, 1948; Sugiura and Stock, 1952) was purchased from CH3 BioSystems and cultured in RPMI-1640-Glutamax medium (Gibco) supplemented with 10% FBS (Eurobio), 10 mM HEPES (Thermo Fisher Scientific), 50 µM β-mercaptoethanol (Gibco) and 100 U/ml penicillin-streptomycin (Thermo Fisher Scientific). The Balb/c mammary carcinoma 4T1 (originating from ATCC and kindly provided by Dr S. Fiorentino) was cultured as described in (Bobrie et al., 2012) in RPMI-1640-Glutamax medium (Gibco) supplemented with 10% FBS (Eurobio), 10 mM Hepes, 100 U/mL penicillin/streptomycin and 1 mM sodium pyruvate (Thermo Fisher Scientific). The MutuDC cell line established from spleen tumors of CD11c:SV40LgT-eGFP-transgenic C57BL/6 mice was kindly provided by Dr. Hans Acha-Orbea (Fuertes Marraco et al., 2012) and cultured in IMDM (Sigma-Aldrich), supplemented with 8% FBS (Biosera), 2mM Glutamax (Gibco), 10 mM HEPES, 50 µM β-mercaptoethanol and 100 U/ml penicillin-streptomycin. The mouse spontaneously immortalized embryonic fibroblast cell line Pfa1 (Seiler et al., 2008) was kindly provided by Dr Sebastian Doll and cultured in DMEM (Thermo Fisher Scientific) supplemented with 10% FBS (Eurobio) and 100 U/ml penicillin-streptomycin. *Mus dunni* and *Mus dunni*-XG7 fibroblasts (CRL-2017) were kindly provided by Dr J.P. Stoye’s lab and cultured in IMDM (Sigma-Aldrich), supplemented with 5% FBS (Eurobio), 2 mM Glutamax (Gibco), 50 µM β-mercaptoethanol and 100 U/ml penicillin-streptomycin. All cell lines were checked for mycoplasma contamination each time a batch was frozen, and found to be negative.

Mouse primary spleen DC were cultured in RPMI-1640-Glutamax medium supplemented with 10% FBS (Biosera), 1% MEM non-essential amino acids (Thermo Fisher Scientific), 1 mM sodium pyruvate, 10 mM HEPES, 50 µM β-mercaptoethanol, and 100 U/ml penicillin-streptomycin.

### Mice

OT1 mice on a Rag2 -/-C57BL/6N background (Lantz et al., 2000) were bred in the CERFE SPF animal facility (Evry, France) for Institut Curie. At 6-16 weeks age, mice were housed in the Institut Curie animal facility for at least 1 week before use. OT1 T cells were obtained by purification with EasySep™ mouse naïve CD8^+^ T cell isolation kit (Stemcell) from spleen and lymph nodes of OT1 mice, according to manufacturer instructions. PanDCs were isolated from spleen of two female C57BL/6 mice aged 14 weeks, using mouse Pan dendritic cell isolation kit according to manufacturer’s instructions (Miltenyi Biotec)(Vremec and Segura, 2013).

### Plasmids and generation of stable EO771 cell lines

The plasmid pmCherry-C1 from Clontech modified to add a myristoylation and palmytoylation sequence plus a linker (ATG**GGCTGCATCAAGAGCAAGCGCAAG**GACAACCTGAACGACGA CGGCGTGGACgaaccggtcgccacc) in Nterm of mCherry was kindly provided by Dr. Franck Perez (Mathieu et al., 2021; Valenzuela and Perez, 2020). myr/palm-mCherry-EO771 cells were sorted based on mCherry expression (S3™ cell sorter, Bio-rad) 3 days after transfection by 5 µl of lipofectamine 2000 (Invitrogen) with 2 µg of the plasmid and regularly re-sorted to maintain expression of myr/palm-mCherry.

For myr/palm-OVA, the pTCP vector (transOMIC) was used as backbone and myristoylation and palmytoylation sequences (same as for mCherry), followed by OVA with in-frame a myc tag, the cleavage peptide P2A and puromycin resistance sequences were included in the coding frame. myr/palm-OVA-EO771 cells were obtained by selection with 4 µg/ml of puromycin, 48 hr after transduction with lentivirus from the supernatant (48 h) of HEK293FT transfected with TransIT-293 and 2 µg of the plasmids (pVSVG, pPAX2 and m/p-OVA, proportions 2:5:8). Stable cells were subsequently cultured in the presence of puromycin 2 µg/ml.

### EVs and ENPs isolation by differential ultracentrifugation and density gradient

To obtain conditioned medium, EO771, Pfa1 and 4T1 cells (plated 1 to 3 days before in complete medium, to reach about 80% confluency) were cultured for 24h in complete medium without serum (= serum-free medium). MutuDC were cultured for 24h in complete medium depleted from serum EV. Cell viability was measured at the end of the culture and found to be more than 85%. EV depletion from serum-containing medium was performed by overnight ultracentrifugation at 100,000x*g* in a 45Ti rotor of complete medium containing 20% FCS as described previously (Thery et al., 2006). Depleted medium was recovered by pipetting, leaving 1cm of medium above the pellet, before 0.22 µm filtration for sterilization (Liao et al., 2019). Conditioned medium was harvested and the isolation protocol was performed in sterile conditions at 4°C. The conditioned medium (90-300 ml) from 100 - 600×10^6^ cells was centrifuged at 350x*g* for 10 min at 4°C. The supernatant was centrifuged at 2,000x*g* for 20 min at 4°C and the 2k pellet was discarded. The supernatant was concentrated with Millipore filters (MWCO = 10kDa, 70 ml) down to 6 ml and ultracentrifuged at 10,000x*g* for 16 min at 4°C in MLA-80 rotor (Beckman Coulter). The pellet was resuspended and washed with PBS at the same conditions (volume, speed and time), as well as for all the washes in this protocol. The obtained pellet was resuspended in PBS at 0.2 µl/10^6^ cells and called 10k. The supernatant of the first 10,000x*g*, depending on the case, was divided in two: one fraction was ultracentrifuged at 200,000x*g* for 50 min at 4°C in MLA-80 rotor to obtain the 200k followed by a wash in the the same volume of PBS and the same centrifugation conditions. The 200k pellet was either resuspended at 1 µl/10^6^ cells of PBS or resuspended in 1 ml of PBS to seed on top of the velocity gradient to obtain the sEVs and VLPs; the other fraction was ultracentrifuged at 200,000x*g* in MLA-80 rotor overnight at 4°C in order to obtain the Mix (washed in 6ml of PBS by overnight centrifugation at 200,000xg and resuspended at 1 µl/10^6^ cells). The velocity gradient was prepared by adding 5 layers of optiprep at decreasing concentrations, bottom to top: 22%, 18%, 14%, 10% and 6% and ultracentrifuged at 187,000x*g* for 1:30h at 4°C in Sw32.1Ti rotor (Beckman Coulter). Afterwards, 8 fractions of 2 ml were collected carefully from top to bottom, washed by adding 4 ml of PBS and centrifuging at 200,000x*g* for 50 min at 4°C, and resuspended at 0.5 µl/10^6^ cells of PBS. The pellets were kept separated or pooled 1-3 (sEVs) and 5-7 (VLPs). The supernatant of the 200,000x*g* for 50 min was ultracentrifuged at 200,000x*g* in MLA-80 rotor overnight at 4°C and washed to obtain the ENPs (finally resuspended at 1 µl/10^6^ cells). Samples were aliquoted and stored at −80°C.

### EVs and ENPs separation by Asymetric Flow Field-Flow Fractionation (AF4)

An AF4 long channel with a frit inlet coupled to the eclipse system (Wyatt Technology, Santa Barbara, CA, USA) was driven by an isocratic pump system including a degasser and an autosampler (Shimadzu, Kyoto, Japan). Detection was performed by an ultraviolet (UV) detector at 280 nm (Shimadzu) and a multi-angle light scattering (MALS) Dawn Helios-II using a laser at 658 nm (Wyatt Technology). The channel was set up with a 350 µm spacer and a 10 kDa regenerated cellulose membrane (Wyatt Technology). PBS supplemented with 0.02% w/v NaN_3_ (TCI chemicals, Tokyo, Japan) and filtered with 0.1 µm polyethersulfone filter (Sigma-Aldrich, Merck Life Science) was used as a mobile phase. All runs were performed at room temperature (20-25 °C).

A detector flow of 1 mL/min was applied in the channel and the sample was injected with an inject flow rate of 0.2 mL/min. An initial cross flow of 2 mL/min for 5 min was applied. Subsequently, the cross flow decreased exponentially from 2 mL/min to 0.1 mL/min over 50 min. The protocol was finished by cleaning the channel from all remaining components using a cross flow of 0 mL/min for 10 min. The elution inject mode was used during the entire run. The Voyager software (Wyatt Technology) was used for data acquisition and the Astra software version 7.3.2 (Wyatt Technology) was used for data analysis. Baseline subtraction was performed. For size distribution analysis the weight density method with results fitting was applied. The sphere model was used. The refractive index for EVs (1.37) was added for correct number density estimation. Light scatter (in relative signal), diameter (in nm) and UV absorption (280nm) were plotted against time.

The AF4 eluted fractions were collected during time and pooled as indicated in Suppl. Fig.2B, and concentrated to 65 µL using Amicon Ultra-2 10K filters (Merck Life Science).

### EV characterization

#### Western Blot

Cell lysates from 2×10^5^ cells, EVs and ENPs secreted by 20×10^6^ cells were resuspended in Laemmli sample buffer (Bio-Rad), boiled 10 min at 95°C and run in 4% - 15% Mini-Protean TGX Stain-Free gels (Bio-Rad) in non-reducing conditions (no β-mercaptoethanol nor DTT). Immuno-Blot PVDF membranes (Bio-Rad) were developed using Immobilon forte western HRP substrate (Millipore). The antibodies used were anti-mouse: CD63 1/200 (clone R5G2, MBL, D263-3), CD9 1/1000 (clone KMC8, BD Bioscience, 553758), Alix 1/1000 (clone 3A9, Cell Signaling, 11/2012), MFGE8 1/1000 (clone 18A2-G10, MBL, D199-3), Hsp90 1/1000 (clone AC88, Enzo Lifescience, ADI-SPA-830-F), Argonaute2 1/1000 (Cell Signaling, 2897), MHC I 1/1000 (rabbit-anti-mouse MHCI, made and kindly provided by Dr H. Ploegh, Boston) (Veron et al., 2005), 14-3-3 (clone EPR6380, Abcam, ab125032), p30 gag MLV 1/1000 (clone R187), env MLV 1/2000 (clone 83A25) these last 2 kindly provided by Dr Leonard Evans.

#### Nanoparticle tracking analysis

Particle concentration, size distribution and zeta potential were measured using ZetaView PMX-120 (Particle Metrix) with software version 8.04.02. Sensitivity was set at 76 and shutter at 70, 11 positions and frame rate at 30. An aliquot of 1 µl from each sample was used for the measurements and dilutions vary between 1/1000 to 1/100000 depending on the concentration of the sample.

#### Cryo-electron microscopy

Cryo-EM was performed on −80°C frozen samples. Lacey carbon 300 mesh grids (Ted Pella, USA) were used in all cryo-EM experiments. Blotting was carried out on the opposite side from the liquid drop and samples were plunge frozen in liquid ethane (EMGP, Leica, Germany). Cryo-EM images were acquired with a Tecnai G2 (Thermo Fisher Scientific, USA) Lab6 microscope operated at 200 kV and equipped with a 4k x 4k CMOS camera (F416, TVIPS). Image acquisition was performed under low dose conditions of 10 e^-^/Å^2^ at a magnification of 50,000 or 6,500 with a pixel size of 0.21 or 1.6 nm, respectively.

#### Protein quantification

Micro BCA™ Protein Assay kit (Thermo Scientific) was used for the protein quantification. An aliquot of 1-5 µl from each sample was used for the measurements and diluted in a final volume of 150 µl.

#### Murine leukemia virus infectivity assay

The infectivity capacity of the MLV-containing samples was assessed as previously described (Young et al., 2012). Briefly, the 200k obtained from 10×10^6^ EO771 or MutuDC cells was added in the presence of polybrene (10 µg/ml) to 3×10^5^ *M*.*dunni* cells transduced with the replication defective plasmid XG7 encoding for GFP (*M*.*dunni-XG7*) in a 6-well plate. The cells were maintained for 14 days in culture, passaged every 2-3 days diluted 1/10-1/25, and production of the XG7 pseudotyped virus was evaluated by incubation of their supernatant with untransduced *M*.*dunni* cells. Briefly, 3×10^3^ *M*.*dunni* cells were incubated with 100 µl of supernatant in the presence of polybrene, and GFP expression was measured by flow cytometry after 3 days.

#### Qualitative and quantitative proteomic analyses

For the qualitative analysis (Fig.1F, Suppl. Table 1), 200k pellets from EO771, 4T1 and 200k + 10k pellets of MutuDC (10 µg, one biological replicate each) were used. For the quantitative analysis (Fig.3, Suppl. Table 2), 5 biological replicates of the 10k, sEVs, VLPs, ENPs and Mix (20 µg each) were used.

##### Sample Preparation

samples were resuspended in 5 µl (qualitative) or 10 µL (quantitative) (2 µg/µL) of 8 M Urea, 200 mM ammonium bicarbonate respectively. After reduction with 5 mM DTT for 30min at 57 °C and alkylation with 10 mM iodoacetamide for 30 min at room temperature in the dark, samples were diluted in 100 mM ammonium bicarbonate to reach a final concentration of 1 M urea. For the qualitative analysis, the 200k pellets were digested for 2 h at 37°C with 0.4 µg of Trypsin/Lys-C (Promega CAT#: V5071) and then overnight by adding 0.4 µg of Trypsin/Lys-C. For quantitative analyses, samples were digested overnight at 37 °C with Trypsin/Lys-C at a ratio of 1/50. Digested samples were loaded onto homemade C18 StageTips for desalting, then eluted using 40/60 MeCN/H_2_O + 0.1 % formic acid and vacuum concentrated to dryness. Peptides were reconstituted in 10 µl of injection buffer in 0.3% trifluoroacetic acid (TFA) before liquid chromatography-tandem mass spectrometry (LC-MS/MS) analysis.

##### LC-MS/MS Analysis

Peptides for the qualitative analysis were separated by reversed phase LC on an RSLCnano system (Ultimate 3000, Thermo Fisher Scientific) coupled online to an Orbitrap Fusion Tribrid mass spectrometer (Thermo Fisher Scientific). Peptides were trapped in a C18 column (75 μm inner diameter × 2 cm; nanoViper Acclaim PepMap 100, Thermo Fisher Scientific) with buffer A (2/98 MeCN:H2O in 0.1% formic acid) at a flow rate of 3.0 µl/min over 4 min to desalt and concentrate the samples. Separation was performed using a 40 cm × 75 μm C18 column (Reprosil C18, 1.9 μm, 120 Å, Pepsep PN: PSC-40-75-1.9-UHP-nC), regulated to a temperature of 40 °C with a linear gradient of 3% to 32% buffer B (100% MeCN in 0.1% formic acid) at a flow rate of 150 nl/min over 91 min. Full-scan MS was acquired using an Orbitrap Analyzer with the resolution set to 120,000, and ions from each full scan were higher-energy C-trap dissociation (HCD) fragmented and analysed in the linear ion trap.

For the quantitative analyses, liquid chromatography (LC) was performed as above with an RSLCnano system (Ultimate 3000, Thermo Scientific) coupled online to an Orbitrap Eclipse mass spectrometer (Thermo Fisher Scientific). Peptides were trapped onto the C18 column at a flow rate of 3.0 µL/min in buffer A for 4 min. Separation was performed on a 50 cm nanoviper column (i.d. 75 μm, C18, Acclaim PepMapTM RSLC, 2 μm, 100Å, Thermo Scientific) regulated to a temperature of 50°C with a linear gradient from 2% to 25% buffer B at a flow rate of 300 nL/min over 91 min. MS1 data were collected in the Orbitrap (120,000 resolution; maximum injection time 60 ms; AGC 4 × 10^5^). Charges states between 2 and 7 were required for MS2 analysis, and a 60 s dynamic exclusion window was used. MS2 scan were performed in the ion trap in rapid mode with HCD fragmentation (isolation window 1.2 Da; NCE 30%; maximum injection time 35 ms; AGC 10^4^).

##### Mass Spectrometry Data analysis

For identification, the data was searched against the *Mus musculus* (UP000000589_10090) UniProt database and a manually curated list of murine virus protein sequences using Sequest-HT through Proteome Discoverer (version 2.4). The database of murine virus proteins includes protein sequences from all known mouse endogenous and exogenous retroviruses (523 sequences manually extracted from Swissprot) and from endogenous MLV envelope glycoproteins (53 sequences), translated from the nucleotide sequences of proviruses annotated as previously described (Attig et al., 2017). Enzyme specificity was set to trypsin and a maximum of two-missed cleavage sites were allowed. Oxidized methionine, Met-loss, Met-loss-Acetyl and N-terminal acetylation were set as variable modifications. Carbamidomethylation of cysteins was set as fixed modification. Maximum allowed mass deviation was set to 10 ppm for monoisotopic precursor ions and 0.6 Da for MS/MS peaks. The resulting files were further processed using myProMS (Poullet et al., 2007) v3.9 (https://github.com/bioinfo-pf-curie/myproms). FDR calculation used Percolator (The et al., 2016) and was set to 1% at the peptide level for the whole study. Proteins were considered expressed if identified with at least 3 peptides among 5 replicates. The label free quantification was performed by peptide Extracted Ion Chromatograms (XICs), reextracted across all conditions and computed with MassChroQ version 2.2.1 (Valot et al., 2011). For protein quantification, XICs from proteotypic and non-proteotypic peptides were used by assigning to the best protein, peptides shared by multiple match groups, and missed cleavages, charge states and sources were allowed. Median and scale normalization was applied on the total signal to correct the XICs for each biological replicate (N=5) for total signal and global variance biases. Label-free quantification (LFQ) was performed following the algorithm as described (Cox et al., 2014) with the minimum number of peptide ratios set to 2 and the large ratios stabilization feature. The final LFQ intensities were used as protein abundance.

For principal component analysis (PCA), data were filtered to only allow proteins with at least 3 quantified peptide ions per sample and with all missing allowed across all samples. The LFQ values of the 3200 proteins selected were log10-transformed and the remaining (27%) missing values were imputed using the R package missMDA (Josse and Husson, 2016).

State-specific protein analysis (SSPA) is an in-house statistical assay that provides statistical ground to the potential state-specificity of a protein (typically present vs absent) based on the distribution of its missing values. To perform this test, peptides XICs are summed for each protein and the resulting value is converted into pseudo-counts by log2-transformation and background subtraction so that the lowest pseudo counts are no smaller than 1. Missing values are converted into 0. For each protein, compared states are then ordered by decreasing mean of their replicates (computed after outlier exclusion) and split into 2 sets (of states) at the largest difference between means of 2 consecutive states (largest step). The protein is declared as overexpressed in the first set and underexpressed in the second. The extent of overexpression is represented by the “best delta” which is the ratio of the largest step over the biggest mean. A statistical test relying on the general linear model with a negative binomial law is also performed to estimate the significance of the difference of means between the 2 sets. Finally, the p-values obtained for the whole dataset are corrected for multiple testing according to the Benjamini-Hochberg (FDR) method.

For heatmap representations, log10-transformed LFQ values for proteins with more than 3 peptides were used. Euclidean distances were calculated for the clustering of proteins (rows) and samples (columns). Pheatmap R package was used for representation. All proteins were used for suppl. Fig4A. The top 15 or 10 most significantly enriched proteins of each sample type (proteins with delta ≥50 ranked by adjusted p-value according to SSPA) were selected for representation in Fig.3D. The 46 quantified viral proteins were used for Fig.3E. Proteins described as DAMPs or PAMPs in (Hoshino et al., 2020) were selected for representation in Fig.4F.

GO-enrichment for Cellular Component was used in two analyses: 1) the top 100 most expressed proteins and 2) the top 100 most SSPA-specific proteins in each group of particles. Enrichr R package was used for enrichment calculations (Kuleshov et al., 2016) and ggplot2 was used for representation. The mass spectrometry proteomics raw data are being deposited to the ProteomeXchange Consortium via the PRIDE (Perez-Riverol et al., 2022) partner repository (dataset identifier number pending).

### mCherry and OVA quantification

The amount of mCherry present in each sample was quantified measuring the fluorescence (excitation 585/emission 625) of 50 µl of each sample through a spectrophotometer (SpectraMax iD3), side-by-side with a standard curve of recombinant mCherry of known concentration. The amount of OVA in each sample was calculated from the signal intensity of the ∼ 40 kDa band, in the Western blot revealed with OVA antibody, using the curve of recombinant OVA with known concentration. Intensity of the bands was quantified using Image Lab software (Bio-rad).

### Uptake, viability and activation assay

MutuDC or isolated panDC were seeded at 1×10^5^ in round bottom 96 well-plates in 50 µl of complete IMDM depleted from serum-derived EVs by overnight ultracentrifugation at 100,000*xg*. Then, 1 or 2 µg of protein (for the phenotypic change assays), or 10 or 30 ng of mCherry (for the uptake assays) of the EVs and ENPs from EO771 m/p-mCherry or controls (medium alone, recombinant mCherry, LPS-EB ultrapure 10 µg/ml (Invivogen) and CpG 2 µg/ml (TriLink) were added in a final volume of 100 µl. After 16h at 37°C, cells were harvested and the supernatant was frozen for later quantification of released cytokines. For MutuDC, cells were stained with Fixable viability dye eFluor™ 780, FcR blocked and stained with fluorochrome-coupled antibodies against mouse: CD40-PerCP-eFluor710, 1/200 (46-0401-80, ThermoFisher), CD86-PE-Cy7, 1/400 (560582, BD Biosciences), PDL1-APC, 1/400 (564715, BD Biosciences) and MHC-II-eFluor 450, 1/200 (48-5320-82, ThermoFisher) or their respective isotypes controls. For panDC, cells were FcR blocked and stained with: B220-FITC, 1/200 (553087, BD Biosciences), CD11c-PerCP-Cy5.5, 1/200 (117328, BioLegend), CD86-PE-Cy7 1/400 (560582, BD Biosciences), XCR1-AF647, 1/200 (148214, BioLegend), MHC-II-APC-Cy7, 1/200 (107628, BioLegend), CD40-BV605, 1/200 (745218, BD Biosciences) and CD172a-BUV737, 1/200 (741819, BD Biosciences) or their respective isotypes controls. Cells were then stained with DAPI. Cells were analyzed with a Cytoflex cytometer (Beckman). PMTs were first set following recommendation of the cytometer, and adjusted if needed to obtain signal at 0 for unstained samples and not saturated in the positive controls. Compensation was performed with single-stained UltraComp eBeads (Thermo Fisher). Data was analyzed with FlowJo software.

### Cytokine bead array

A customized set of 17 cytokines (IFNγ, IL2, IL5, IL4, IL6, MCP1 (CCL2), IL13, IL10, IL17a, MIP1γ, TNFa, MIP1β, IL12 p70, RANTES (CCL5), MIG (CXCL9), IL1α, IL1β) reported to be secreted by DCs (Morelli et al., 2001) was selected to measure in the supernatant of the MutuDC by Cytometric Bead Array (BD CBA Flex Sets) following the manufacturer’s recommendations. 15 µl of each culture supernatant was used for the assay. Samples were acquired on a BD FACS verse cytometer and analyzed with the FCAP Array software.

### Cross-presentation assay

MutuDC were seeded at 1×10^4^ cells in round bottom 96 well-plates with complete EV-depleted RPMI and incubated for 5h with 1 or 2 ng of OVA in the EVs and ENPs or controls (medium alone, recombinant OVA and SIINFEKL peptide curves). After what, they were washed once with 37°C RPMI and cocultured with 5×10^4^ OT1 T cells obtained by purification with EasySep™ mouse naïve CD8^+^ T cell isolation kit (Stemcell) from spleen and lymph nodes of OT1 mice. After 16h cells were harvested and activation of the OT1 was measured by CD69 (553237, BD Pharmingen) and CD25 (551071, BD Biosciences) expression. Cells were also stained with Vα2 TCR (560624, BD Pharmingen) and CD8a (553035, BD Pharmingen) for identification. Cells were analyzed with a Cytoflex cytometer (Beckman). Data was analyzed with FlowJo software.

## ACKNOWLEDGMENTS

We thank for fruitful discussions and tools: Drs J. Helft, L. Saveanu, H. Acha-Orbea, L. Evans, J.P. Stoye, G. Boncompain, S. Doll, S. Fiorentino, L. Alaoui, M. Burbage, M. Gros, M. Jouve-San Roman, M. Rehmsmeier, L. Ringrose, and C. Basse.

This work was funded by INSERM, CNRS, Institut Curie, French IdEx and LabEx (ANR-10-IDEX-0001-02 PSL), the MSCA-ITN grant “TRAIN-EV” agreement No 722148, grants from french ANR (ANR-18-CE13-0017-03; ANR-18-CE15-0008-01; ANR-18-CE16-0022-02), INCa (INCa_16083), Fondation ARC (PGA12021020003189_3588), FRM (FDT202106013265, EQU201903007925 and DGE20121125630), Cancéropôle Île-de-France (2013-2-EML-02-ICR-1), financial support from “la Région Île-de-France” (N°EX061034) and ITMO Cancer of Aviesan and INCa on funds administered by Inserm (N°21CQ016-00) for MS analysis. We also acknowledge the following Core facilities of Institut Curie: Cell and Tissue Imaging (PICT-IBiSA), member of the French national research infrastructure France-BioImaging (ANR10-INBS-04) for fluorescence and electron microscopy, Cytometry for assistance in data acquisition, Extracellular Vesicles for assistance in EV isolation and NTA-based quantification.

## EXPANDED VIEW FIGURE LEGENDS

**Suppl. Fig.1:**
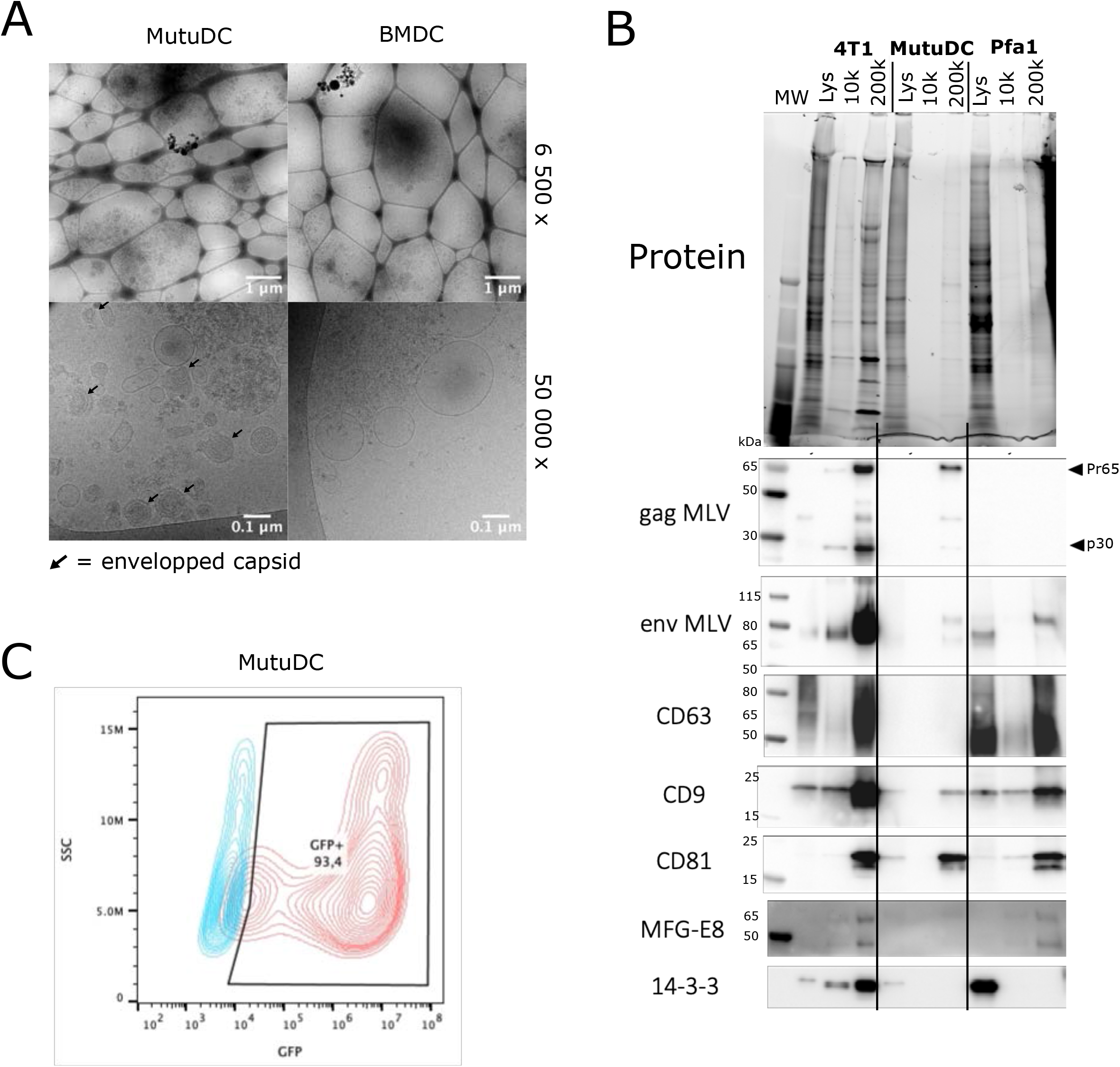
Characterization of the EVs from 4T1 and MutuDC cell lines, primary bone marrow-derived DCs (BMDC), and immortalized fibroblasts Pfa1. (A) Cryo-EM of 200k from MutuDC and primary BMDCs, showing presence of enveloped capsids in the former (arrows). (B) Western blot of cell lysate (Lys), 10k and 200k of 4T1, MutuDC and Pfa1, showing total proteins (top) and hybridization with antibodies against viral and endogenous proteins as indicated (bottom). Gag and env are detected in pellets from 4T1 and MutuDC, and env is detected in Pfa1. (C) infectivity assay performed as in Fig.1H, with the 200k pellets of MutuDC, showing presence of infectious virus.

**Suppl. Fig.2:**
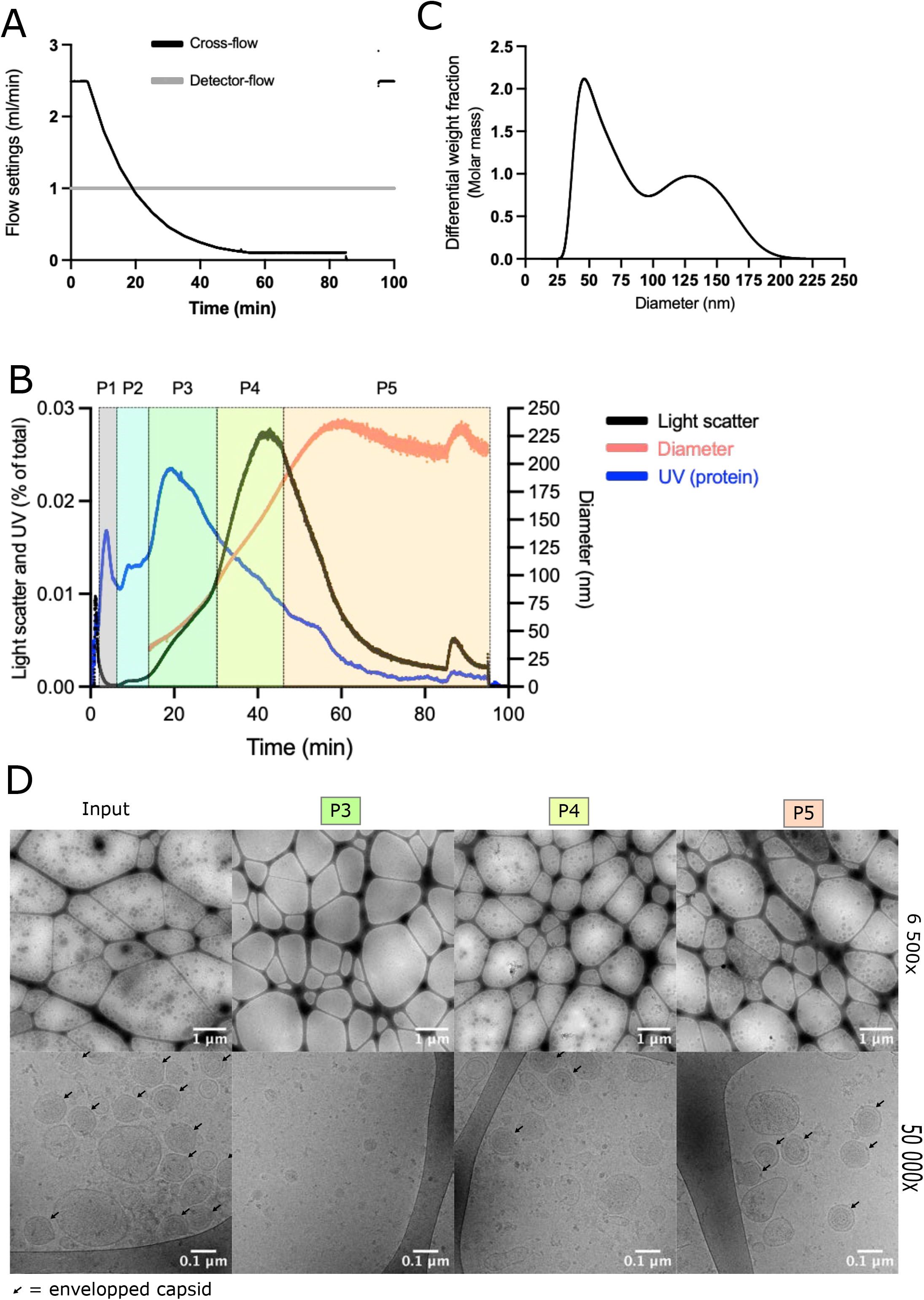
(A) Detector-flow and cross-flow settings used for the AF4-based separation of EVs from a EO771 200k pellet. (B) UV (280 nm), Light Scattering, and particle size measured in the fractions recovered overtime (one experiment). UV signal in the absence of light scatter signal and of detected particles ≥ 35 nm (P2) corresponds to non-vesicular proteins or particles. (C) Differential weight fraction calculated from the UV (280 nm) absorption, as a function of the diameter. Two peaks of mass are observed, suggesting the presence of 2 populations of particles of around 35-80 nm and 80-180 nm, named P3 and P4. (D) cryo-EM of the total input and P3, P4 and P5 fractions of B, showing the presence of VLPs in both P4 and P5, together with EVs.

**Suppl. Fig.3:**
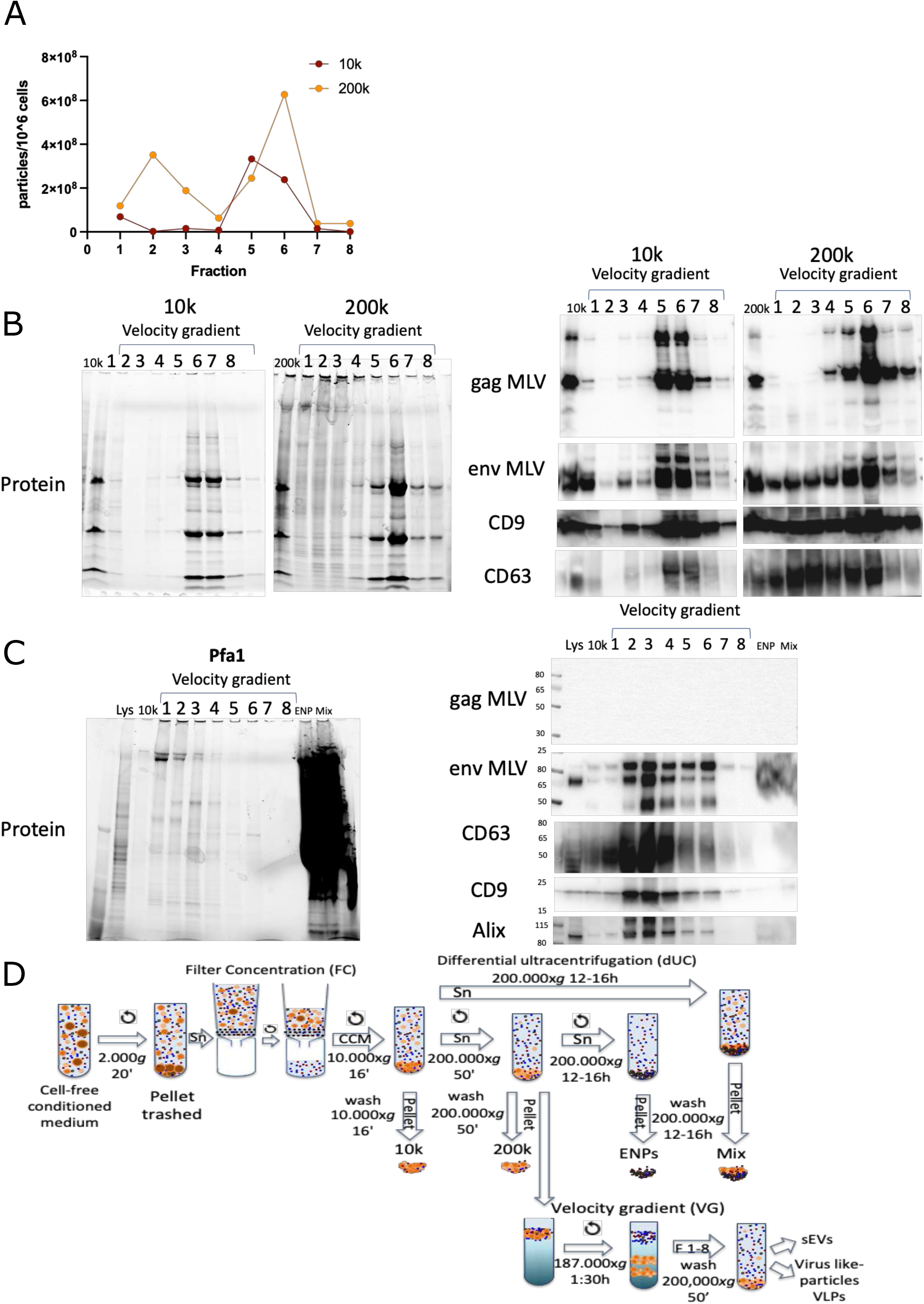
(A) Quantification of particles in fractions recovered from an iodixanol velocity gradient top-loaded with a EO771 10k or 200k pellet from the same CM. A single peak of particles recovered in fractions 5-6 is obtained from the 10k (B) Western blot analysis of the velocity gradient fractions. Stain-free total protein (left) and hybridization with antibodies against gag and env MLV proteins or CD9 and CD63 tetraspanins (right). (C) Western blot analysis of the velocity gradient fractions of 200k obtained from Pfa1. Stain-free total protein stain (left) and hybridization with antibodies against gag and env MLV proteins or CD9, CD63 and Alix EV proteins (right). (D) Scheme of the final combined protocol used to recover 10k, sEVs, VLPs, ENPs, 200k and Mix.

**Suppl. Fig.4.**
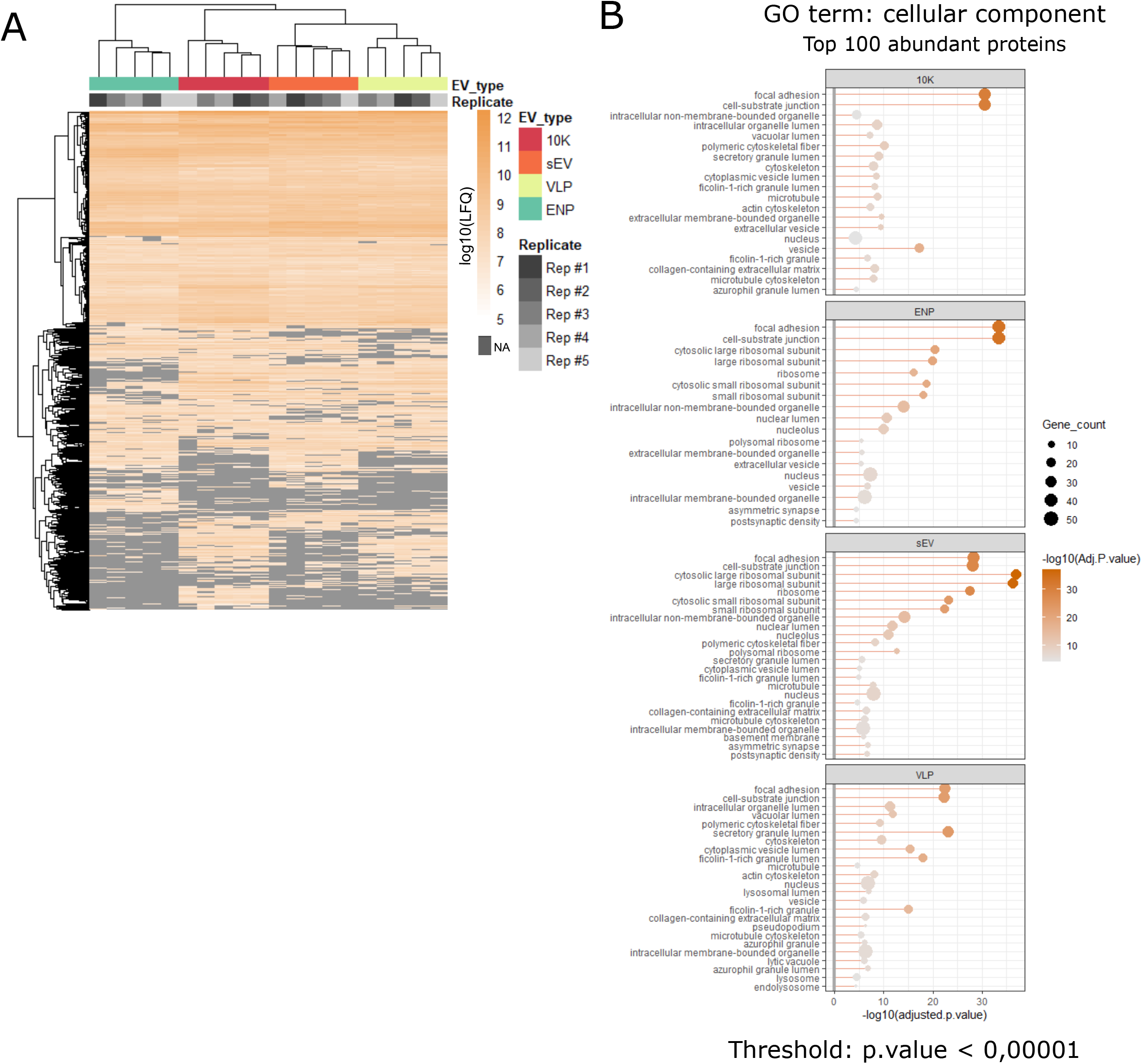
Label-free quantification analysis of the proteomic results. (A) Heatmap of the protein abundance of all proteins identified in one of the 5 groups. NA= not detected/absent. (B) GO-term enrichment analysis of the top-100 most abundant proteins (according to LFQ) in the 4 groups. GO-terms with p<0.0001 are represented. GO-terms are ordered by adjusted.p-value.

**Suppl. Fig.5:**
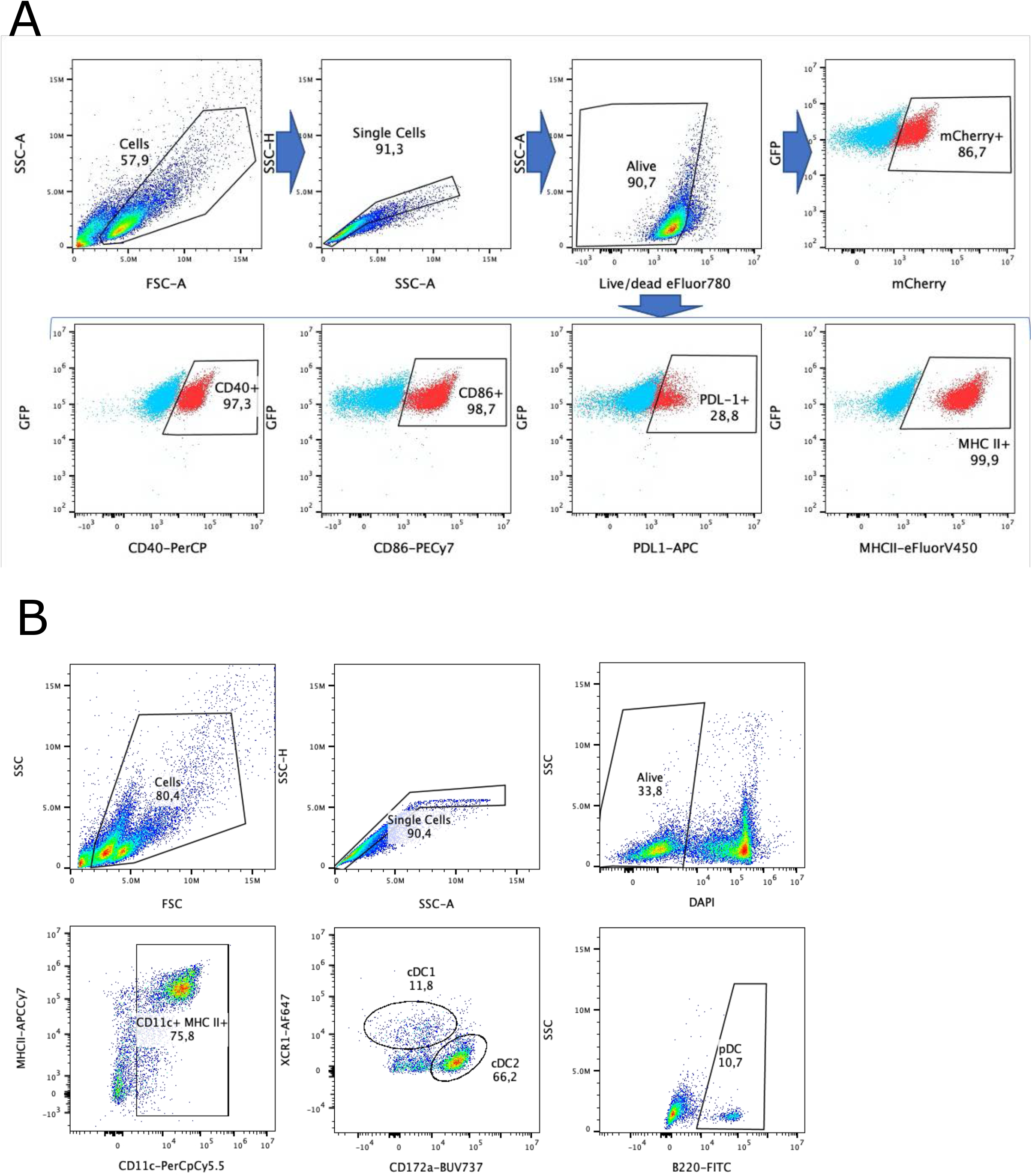
Flow cytometry gating strategy for analysis of dendritic cells’ phenotype upon exposure to EO771 EVs/ENPs. (A) Gating strategy applied to MutuDCs to quantify viability and cell surface markers (shown in Fig.4), and mCherry uptake (shown in Fig.5). A representative example of MutuDCs exposed to 10k is shown. Blue = isotype control, red = specific antibody. (B) Gating strategy applied to spleen DCs (CD11c+) to identify DC subpopulations (cDC1 = XCR1+, cDC2 = CD172+, pDC = B220+) and viability (DAPI-).

***Suppl. Table 1:***Proteomic identification of mouse viral proteins in EV preparations from EO771 (200K pellet), 4T1 (200k pellet) and MutuDCs (10k + 200k mixed pellet) cells. Number of peptides identified in each sample (Tab: results), full list of identified proteins (Tab: all), and list of proteins identified by proteotypic peptides or as endogenous retrovirus proteins (Tab: proteotypic or endogenous, displayed in Fig. 1F) are provided.

***Suppl. Table 2.***Proteomic analysis results. Peptide numbers for all identified proteins, used to generate Fig.3A, are shown in “Venn Diagram” tab. Label-free quantification of each protein is shown in “LFQ quant” tab. State-specific protein analysis (SSPA) results, to identify proteins specifically enriched in one or more sample subtypes, are shown in”SSPA” tab.

